# The Multivariate Genome-wide Architecture of Interrelated Literacy, Language and Working Memory Skills Reveals Distinct Etiologies

**DOI:** 10.1101/2020.08.14.251199

**Authors:** Chin Yang Shapland, Ellen Verhoef, George Davey Smith, Simon E. Fisher, Brad Verhulst, Philip S. Dale, Beate St Pourcain

## Abstract

There is genetic overlap between many measures of literacy, language and phonological working memory (PWM) though our knowledge of multivariate genetic architectures is incomplete. Here, we directly modeled genetic trait interrelationships in unrelated UK youth (8-13 years, N=6,453), as captured by genome-wide relationship matrices, using novel structural equation modeling techniques. We identified, besides shared genetic factors across different domains (explaining 91-97% genetic variance in literacy-related measures such as passage reading fluency, spelling, phonemic awareness, 44% in oral language and 53% in PWM), evidence for distinct cognitive abilities; trait-specific genetic influences ranged between 47% for PWM to 56% for oral language. Among reading fluency measures (non-word, word and passage reading), single-word reading was genetically most diverse. Multivariate genetic and residual covariance patterns showed concordant effect directionality, except for near-zero residual correlations between oral language and literacy-related abilities. These findings suggest differences in etiological mechanisms and trait modifiability even among genetically highly correlated skills.

## Introduction

Within most Indo-European languages, including English, an alphabetic writing system maps sequences of symbols to the sounds and meaning of words [1], where symbols or graphemes (letters or groups of letters) represent individual sounds (phonemes). This correspondence of spoken language to printed words, known as the alphabetic principle [1], is, in turn, the basis of phonological decoding skills, which enable the interpretation of texts through phonetic transformation [2]. As a consequence, close and reciprocal interrelationships manifest between phonological decoding, letter knowledge, and phonological awareness [1], the ability to dissect and transform words according to their phonological structure [3]. As these links emerge and can be used in real-time to identify printed words, children apply this knowledge to develop reading and spelling skills [1]. Once mastered, reading has a profound impact on the acquisition of knowledge, including print exposure [4], and thus on final educational attainment [5].

Based on this framework, the “Simple View of Reading” [6, 7] proposed that reading comprehension is the product of two independent skill sets, “decoding” and “language comprehension”, where the latter specifically refers to listening comprehension [6], i.e. the ability to interpret the meaning of words in grammatical structures (sentences) presented orally [6]. In younger children, reading comprehension is thought to be constrained primarily by decoding abilities, whereas in older children, with a high level of decoding ability, reading comprehension is mainly a function of language comprehension [8]. The Reading Systems Framework [9] also implicates other general cognitive resources in reading processes, such as phonological working memory (PWM) [10], which is typically assessed by non-word repetition tasks [11]. This task requires a purely sound-based division of non-words along with retention and later reproduction of the sound sequence [10]. Like phonological awareness, PWM has been hypothesized to contribute to decoding.

Recent meta-analytic structural equation modeling studies have reported strong to moderate interrelations between language, literacy and related traits, such as working memory, spanning childhood and adolescence [12]. A considerable part of these relationships can be attributed to shared genetic factors as shown by twin research [13, 14] and by studies of unrelated individuals using genome-wide genotyping information [15]. These findings add to the widely established evidence that individual language- and literacy-related abilities have moderate to strong heritability from mid-childhood onwards [13-26]. Such traits include reading decoding, fluency and comprehension [13, 16-20], spelling [14, 17], phonological awareness [21], language comprehension [26], non-word repetition (a proxy of PWM) [26]. Within twin studies, strong genetic correlations have been identified between word reading efficiency and both reading comprehension and spelling [14, 17, 18]. Moderate to strong genetic links have been found between oral language and reading comprehension [13, 14] and moderate genetic overlap between oral language and word reading efficiency [13, 14]. Within studies investigating common DNA markers in population-based samples, too, modest to moderate genetic correlations have been reported between multiple language- and literacy-related abilities [15].

While there is ample support for shared overarching genetic factors [10, 11, 19-22], we lack a comprehensive map of the genetic architecture which includes both these broad connections as well as unique genetic relationships that may, for example, exist for meaning-based (i.e. comprehension-related) versus code-based (i.e. decoding-related) aspects of language- and literacy-related ability [10].

Thus far, knowledge of multivariate genetic architectures across language and literacy has been largely based on twin analyses [13-15, 17, 18]. Here, we investigate multivariate genetic architectures as directly captured by genome-wide information (using genetic relationship-matrices, GRMs) assessed in unrelated youth from The Avon Longitudinal Study of Parents and Children (ALSPAC) birth cohort [27]. We apply GRM structural equation modeling (GSEM) [27], informed by twin research-modeling techniques, dissecting phenotypic variation into additive genetic and residual variance. To facilitate the interpretability of shared genetic factor structures without loss of model fit in residual variance structures, we define a combined Independent Pathway / Cholesky (IPC) model. This model structures the genetic variance as an Independent Pathway model (with shared and specific genetic factors), while decomposing the residual variance according to a Cholesky model.

Our overarching goal is to understand the relation of reading skills to abilities that are outside of reading proper, but are nonetheless literacy-related. Specifically, we model the multivariate polygenetic architecture across three phenotypic domains: literacy-related measures (reading fluency, spelling, phonemic awareness), oral language (listening comprehension) and PWM (non-word repetition). We, eventually, compare genetic and residual architectures as the developmental relationship between language, phonological decoding and subsequent reading and spelling abilities [1, 2, 6, 7] may implicate additional etiological factors. These additional influences are not captured by genotyped markers involving rare, non-additive or un-tagged genetic influences [28], but also environmental risk factors, as well as random error.

## Results

### Univariate and bivariate genetic architecture of literacy-, language-, and PWM-related abilities

Variability in literacy-, language-, and PWM-related abilities during mid-childhood and adolescence (Table 1) was moderately heritable, confirming previous reports [15, 29]. GCTA SNP-based heritability (SNP-h^2^) estimates for reading (non-word-reading speed and accuracy, word reading speed and accuracy, passage reading speed and accuracy), spelling accuracy, phonemic awareness, listening comprehension and non-word repetition ranged between 30% (SE=0.06) and 50% (SE=0.07) when assessed in unrelated children and adolescents from the general population, irrespective of phenotypic transformation (Supplementary Table 1). All scores were phenotypically (Supplementary Table 2) and genetically (Supplementary Table 3 and 4) correlated (Supplementary Figure 1) ranging between 0.2 to 0.8 and 0.4 to 0.98, respectively.

**Table 1:**
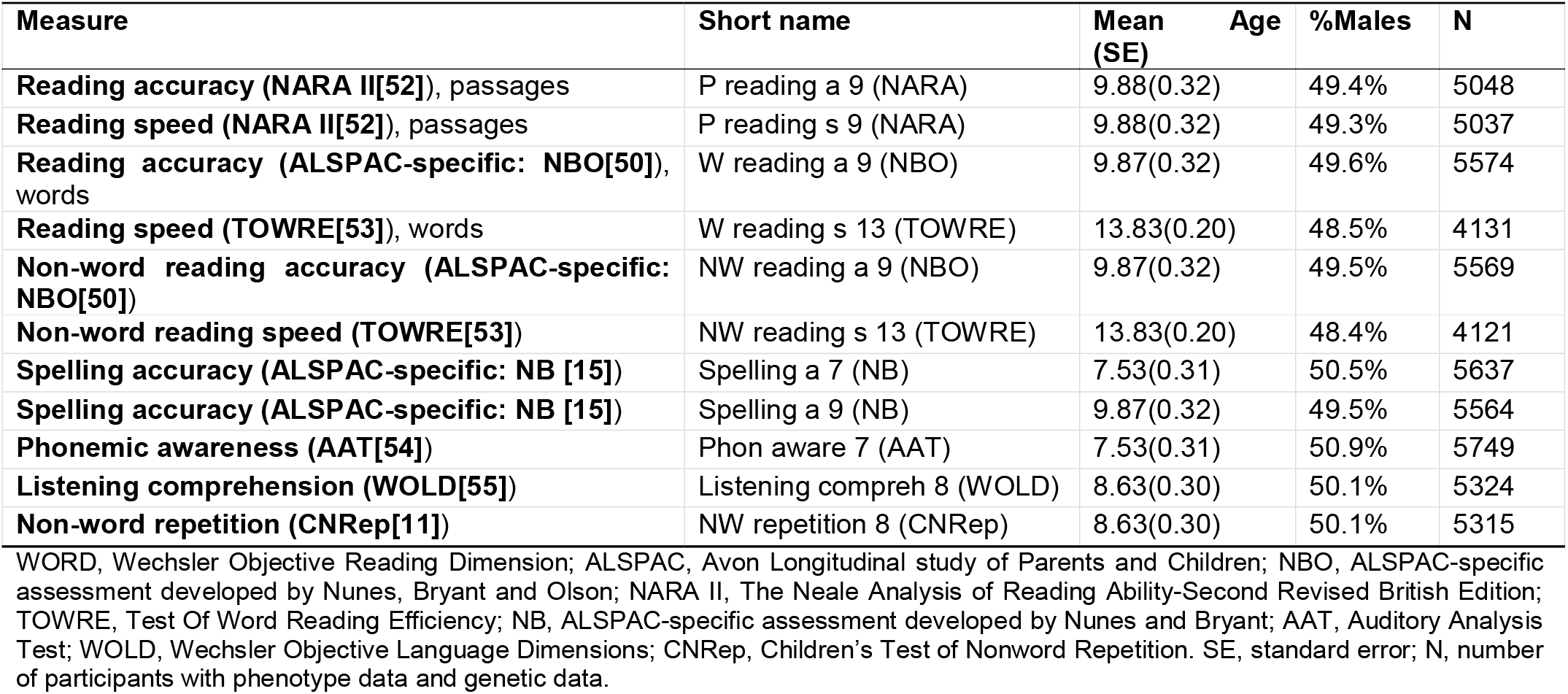
Description of literacy-, language-, and PWM-related abilities

**Table 2:**
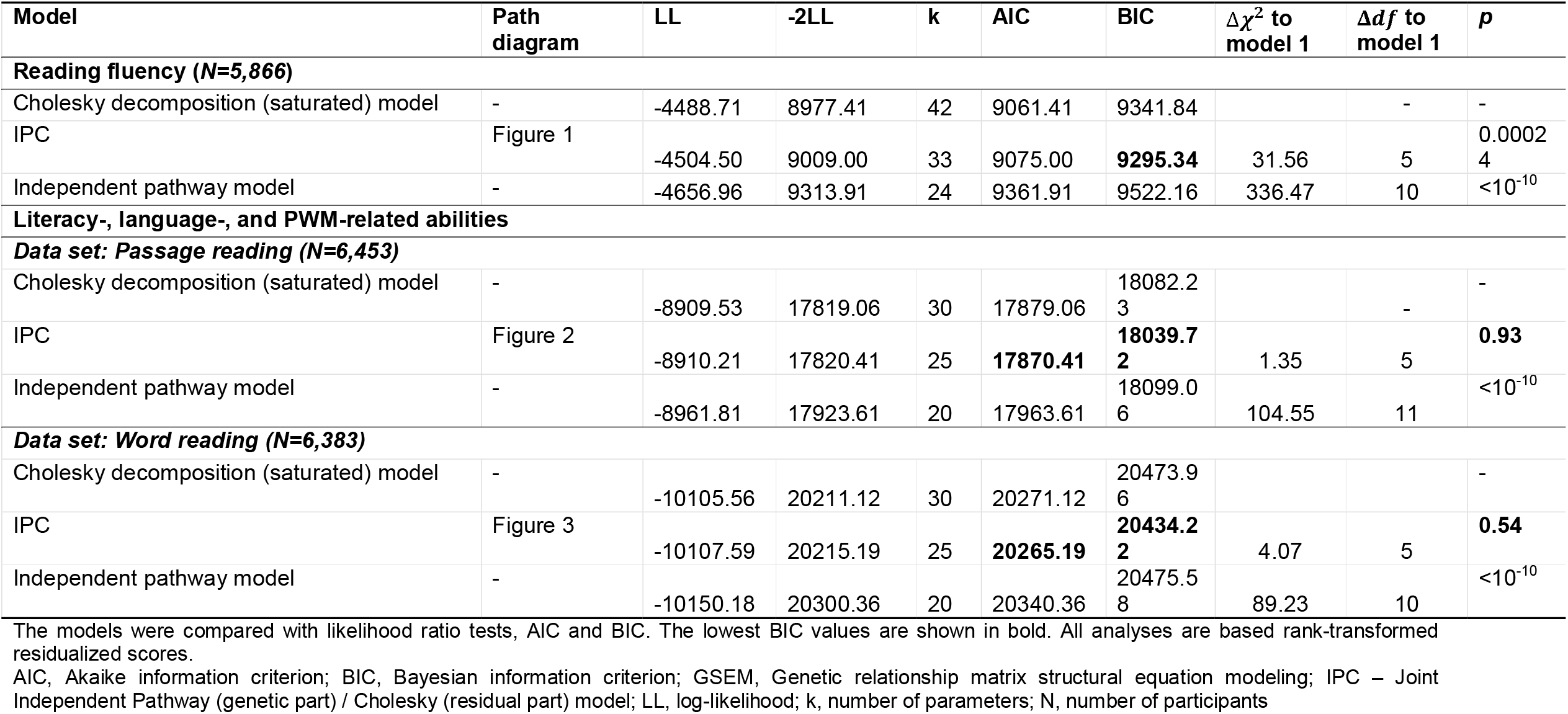
GSEM model fit comparisons

| Model | Path diagram | LL | -2LL | k | AIC | BIC | $\Delta\chi^2$ to model 1 | $\Delta df$ to model 1 | p |
| --- | --- | --- | --- | --- | --- | --- | --- | --- | --- |
| **Reading fluency (N=5,866)** |  |  |  |  |  |  |  |  |  |
| Cholesky decomposition (saturated) model | - | -4488.71 | 8977.41 | 42 | 9061.41 | 9341.84 |  | - | - |
| IPC | Figure 1 | -4504.50 | 9009.00 | 33 | 9075.00 | **9295.34** | 31.56 | 5 | 0.00024 |
| Independent pathway model | - | -4656.96 | 9313.91 | 24 | 9361.91 | 9522.16 | 336.47 | 10 | <10<sup>-10</sup> |
| **Literacy-, language-, and PWM-related abilities** |  |  |  |  |  |  |  |  |  |
| **Data set: Passage reading (N=6,453)** |  |  |  |  |  |  |  |  |  |
| Cholesky decomposition (saturated) model | - | -8909.53 | 17819.06 | 30 | 17879.06 | 18082.23 |  | - | - |
| IPC | Figure 2 | -8910.21 | 17820.41 | 25 | **17870.41** | **18039.72** | 1.35 | 5 | **0.93** |
| Independent pathway model | - | -8961.81 | 17923.61 | 20 | 17963.61 | 18099.06 | 104.55 | 11 | <10<sup>-10</sup> |
| **Data set: Word reading (N=6,383)** |  |  |  |  |  |  |  |  |  |
| Cholesky decomposition (saturated) model | - | -10105.56 | 20211.12 | 30 | 20271.12 | 20473.96 |  |  | - |
| IPC | Figure 3 | -10107.59 | 20215.19 | 25 | **20265.19** | **20434.22** | 4.07 | 5 | **0.54** |
| Independent pathway model | - | -10150.18 | 20300.36 | 20 | 20340.36 | 20475.58 | 89.23 | 10 | <10<sup>-10</sup> |

The models were compared with likelihood ratio tests, AIC and BIC. The lowest BIC values are shown in bold. All analyses are based rank-transformed residualized scores.
AIC, Akaike information criterion; BIC, Bayesian information criterion; GSEM, Genetic relationship matrix structural equation modeling; IPC – Joint Independent Pathway (genetic part) / Cholesky (residual part) model; LL, log-likelihood; k, number of parameters; N, number of participants

**Figure 1:**
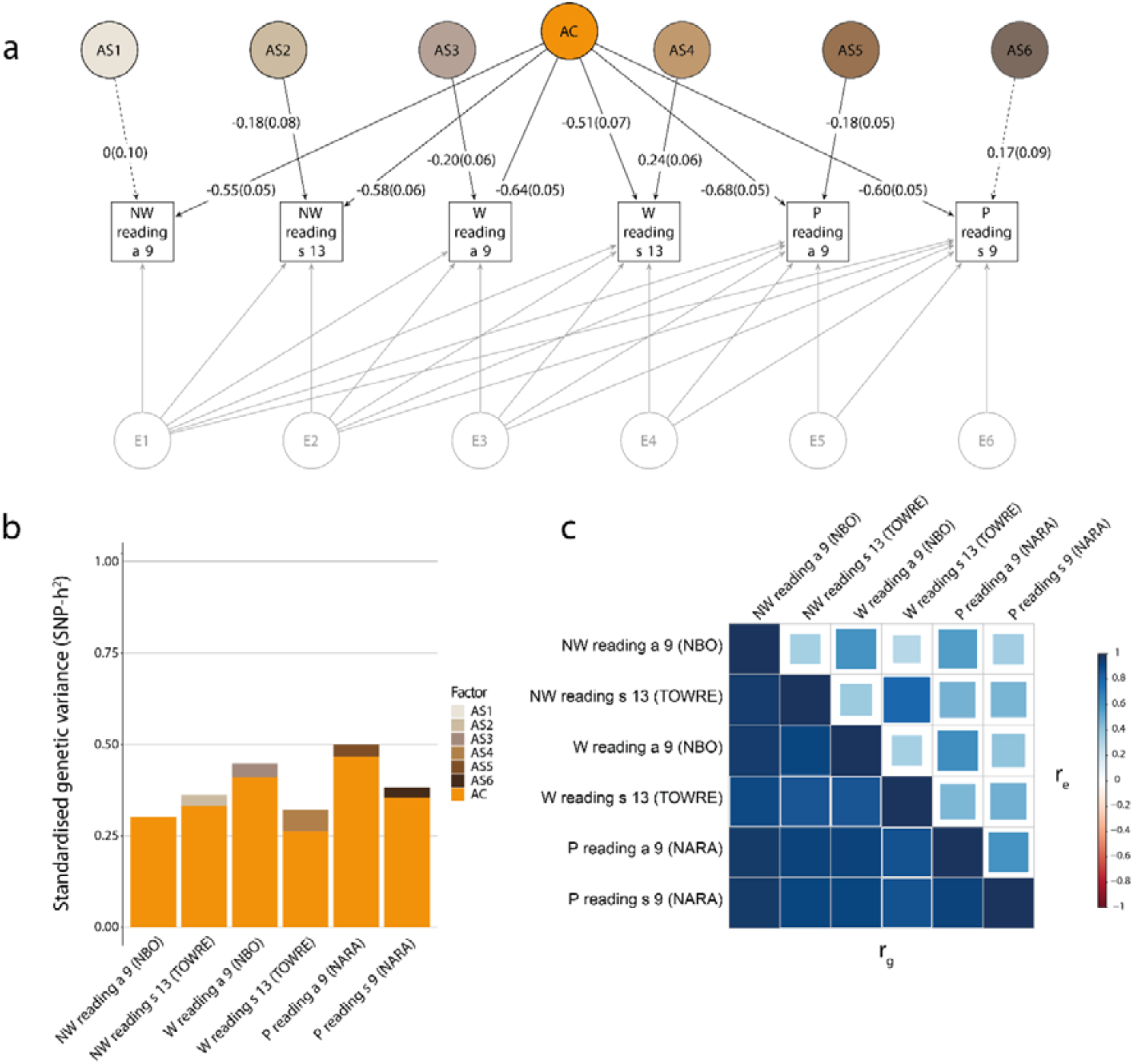
Structural model of reading fluency. Genetic-relationship matrix structural equation modeling (GSEM) of six reading measures (N= 5,866). The path diagram **(a)** depicts the genetic factors of the best-fitting model (IPC model) describing variation in non-word reading accuracy at age 9 (NW reading a 9, NBO), non-word reading speed at age 13 (W reading s 13, TOWRE), word reading accuracy at age 9 (W reading a 9, NBO), word reading speed at age 13 (W reading s 13, TOWRE), passage reading accuracy at age 9 (P reading a 9, NARA) and passage reading speed at age 9 (P reading s 9, NARA). The phenotypic variance was dissected into latent common (AC) and specific (AS1-AS6) genetic factors, according to an Independent Pathway model, as well as latent (E1-E6) residual factors, based on a Cholesky decomposition (not shown, Supplementary Table 6, Supplementary Figure 3). Observed measures are represented by squares and latent factors by circles. Single headed arrows (paths) define relationships between variables. Dotted and solid paths represent factor loadings with *p*>0.05 and *p*≤0.05 respectively. The variance of latent variables is constrained to unit variance; this is omitted from the diagram to improve clarity. The variance plot **(b)** depicts the standardized genetic variance components for the model in (a). The correlogram **(c)** shows genetic (r_g_, lower triangle) and residual (r_e_, upper triangle) correlations for the model in (a). Point estimates and their SEs are reported in Supplementary Table 9.

### Modeling strategy for multivariate analyses

Using a series of structural equation models, we describe the multivariate polygenic genetic architecture of literacy-, language-, and PWM-related abilities. Due to the computational burden of multivariate genetic variance analyses [27], we were not able to conduct one large combined model of all 11 measures in this study. Instead, we adopted a two-stage approach: First, we conducted multiple smaller-scale multivariate models focussed on literacy measures only with the aim of identifying proxy measures that sufficiently capture the observed genetic structures of reading fluency (6 measures) and spelling (2 measures), which were assessed with multiple psychological instruments (Stage 1, Supplementary Table 5). Second, we fitted multivariate genetic models across traits including proxy measures of reading fluency and spelling (identified from Stage 1) as well as other literacy-related measures (phonemic awareness), listening comprehension and non-word repetition (Stage 2, Supplementary Table 5).

For each fitted multivariate model during either stage (except for spelling), three GSEM submodels were examined: (i) a saturated model (Cholesky decomposition), (ii) an independent pathway model and (iii) an IPC model, involving a pathway model with a mixed structure, with an independent pathway model (IP) for the genetic part and a Cholesky decomposition (C) model for the residual part of the model. The model fit was compared using Akaike Information Criterion (AIC) and Bayesian Information Criterion (BIC) fit indices and likelihood ratio tests (LRT). The two spelling measures (Stage 1) were fitted with a Cholesky decomposition model.

### Structural models of reading fluency

We started the process of proxy measure identification (Stage 1) by structurally modeling six reading fluency measures (Supplementary Table 5): non-word reading accuracy and speed, word reading speed and accuracy, and passage reading accuracy and speed, all ascertained in ALSPAC participants between the ages of 9 to 13 years (Table 1, Figure 1). Following BIC (the more stringent criterion), the most parsimonious model describing the data was the IPC model (Figure 1, Table 2, Supplementary Table 6-9). The model identified evidence for a shared latent genetic factor (A) across the six reading fluency measures (Figure 1), in addition to specific genetic factors for some of the measures.

For the shared genetic factor (Figure 1a,b), the strongest factor loading was estimated for passage reading accuracy, explaining 47%(SE=0.07) of the phenotypic variation (Supplementary Table 6) and 93%(SE=4%) of the SNP-h^2^ (Supplementary Table 7). The weakest factor loading was found for word reading speed (Figure 1a,b), explaining only 27%(SE=0.07) of the phenotypic variation (Supplementary Table 6), but 82%(SE=9%) of the SNP-h^2^ (Supplementary Table 7). Thus, the shared genetic factor captured the majority of genetic variance across all reading fluency measures, and its factor structure is reflected by near-perfect genetic correlations (r_g_) between the different reading fluency measures (Supplementary Table 8, Figure 1c). For example, passage reading accuracy had r_g_ of 0.97(SE=0.02) with non-word reading accuracy, 0.92(SE=0.04) with non-word reading speed, 0.92(SE=0.03) with word reading accuracy, 0.88(SE=0.05) with word reading speed, and 0.93(SE=0.04) with passage reading speed.

There was further evidence for trait-specific factors describing additional genetic variation underlying non-word reading speed (passing the nominal *p*-value threshold only), word reading speed and accuracy, and passage reading accuracy (Figure 1a), although the explained phenotypic variation was small. Word reading speed (age 13) had the strongest trait-specific factor loading explaining 6%(SE=0.03) of phenotypic variance (Supplementary Table 6), corresponding to 18%(SE=0.09) of the SNP-h^2^ (Supplementary Table 7)(Figure 1b).

To ensure that the conclusion from Stage 2 analysis would be independent of the choice of the reading fluency measure, we selected two proxy measures based on the structural model for reading fluency in Stage 1: (i) passage reading accuracy (age 9), because it showed the highest common factor loading for reading, and (ii) word reading speed (age 13), because it showed the highest trait-specific genetic factor loading.

### Structural models of spelling

As a further proxy identification model in Stage 1, we fitted a saturated Cholesky model to spelling accuracy scores (Supplementary Table 5) at 7 and 9 years of age (Supplementary Figure 2, Supplementary Table 10-12). This model showed that genetic factors underlying spelling accuracy at age 7 years can capture nearly all (94% with SE=0.09) of the genetic variation for spelling accuracy at age 9 years (Supplementary Table 11), with r_g_ of 0.97 (SE=0.05, Supplementary Table 12). Consequently, the former was selected as a proxy for Stage 2 analysis.

**Figure 2:**
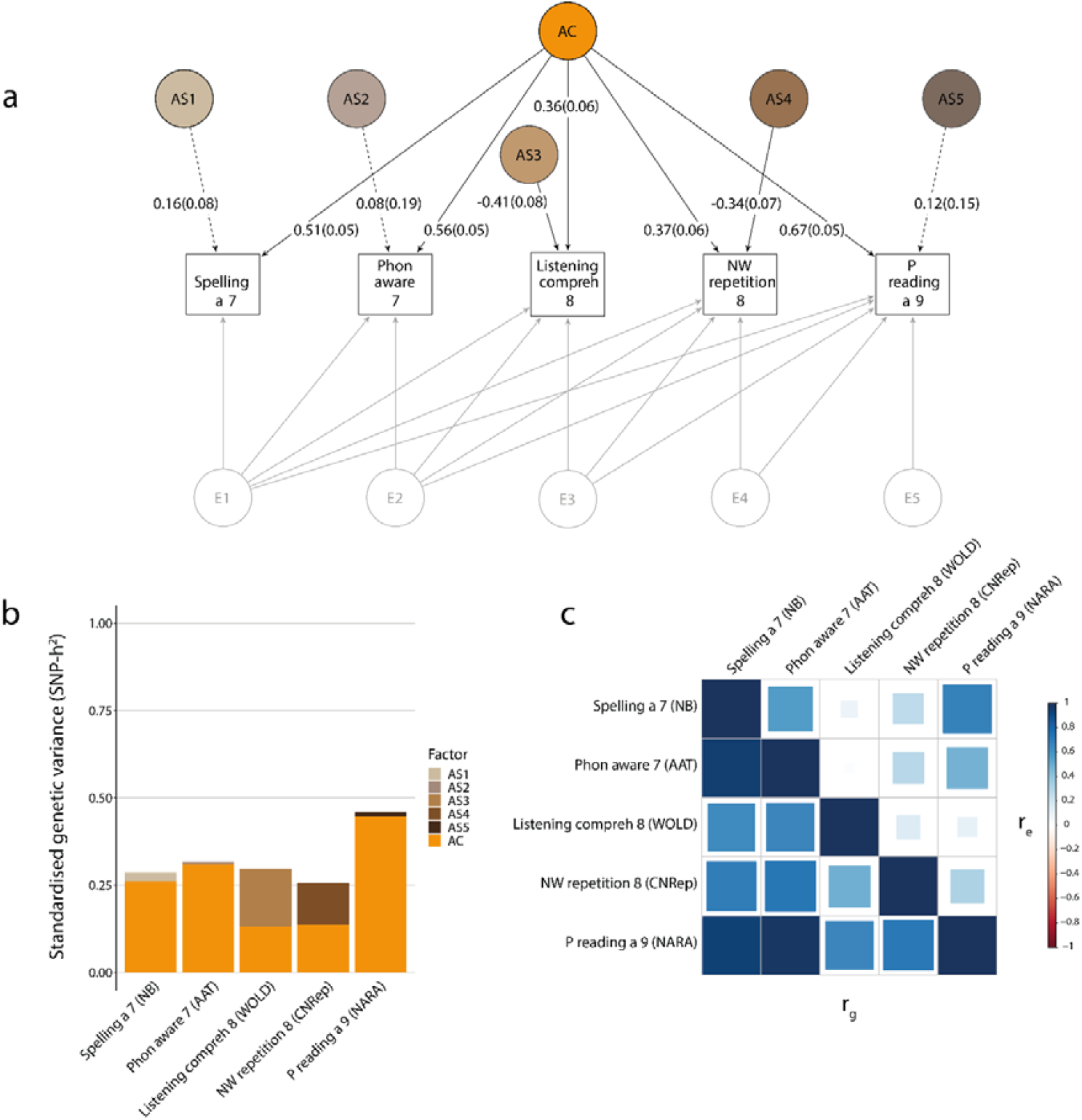
Structural model of literacy-, language-, and PWM-related abilities (Data set: passage reading) Genetic-relationship matrix structural equation modeling (GSEM) of five literacy-, language-, and PWM-related abilities) (N=6,453). The path diagram **(a)** depicts the genetic factors of the best-fitting model (IPC model) describing variation in spelling accuracy at age 7 (Spelling a 7, NB), phonemic awareness at age 7 (Phon aware 7, AAT), listening comprehension at age 8 (Listening compreh 8, WOLD), non-word repetition at age 8 (NW repetition 8, CNRep) and passage reading accuracy at age 9 (P reading a 9, NARA). The phenotypic variance was dissected into latent common (AC) and specific (AS1-AS5) genetic factors, according to an Independent Pathway model, as well as latent (E1-E5) residual factors, based on a Cholesky decomposition (not shown, Supplementary Table 13, Supplementary Figure 4). Observed measures are represented by squares and latent factors by circles. Single headed arrows (paths) define relationships between variables. Dotted and solid paths represent factor loadings with *p*>0.05 and *p*≤0.05 respectively. The variance of latent variables is constrained to unit variance; this is omitted from the diagram to improve clarity. The variance plot **(b)** depicts the standardized genetic variance components for the model in (a). The correlogram **(c)** shows genetic (r_g_, lower triangle) and residual (r_e_, upper triangle) correlations for the model in (a). Point estimates and their SEs are reported in Supplementary Table 16.

### Structural models of literacy-, language-, and PWM-related abilities

For Stage 2 analyses, we investigated the multivariate genetic architecture of literacy-related skills (spelling, phonemic awareness, reading fluency), language-related ability (listening comprehension) and PWM (non-word repetition). Each of the two selected proxy measures for reading fluency was independently studied. The subset including passage reading accuracy (age 9) was referred to as passage reading dataset (Figure 2), while the subset including word reading speed (age 13) was referred to as word reading dataset (Figure 3). For each of these two selected datasets, we compared the fit of a saturated Cholesky decomposition to the fit of an independent pathway and IPC model, as shown in Table 2.

**Figure 3:**
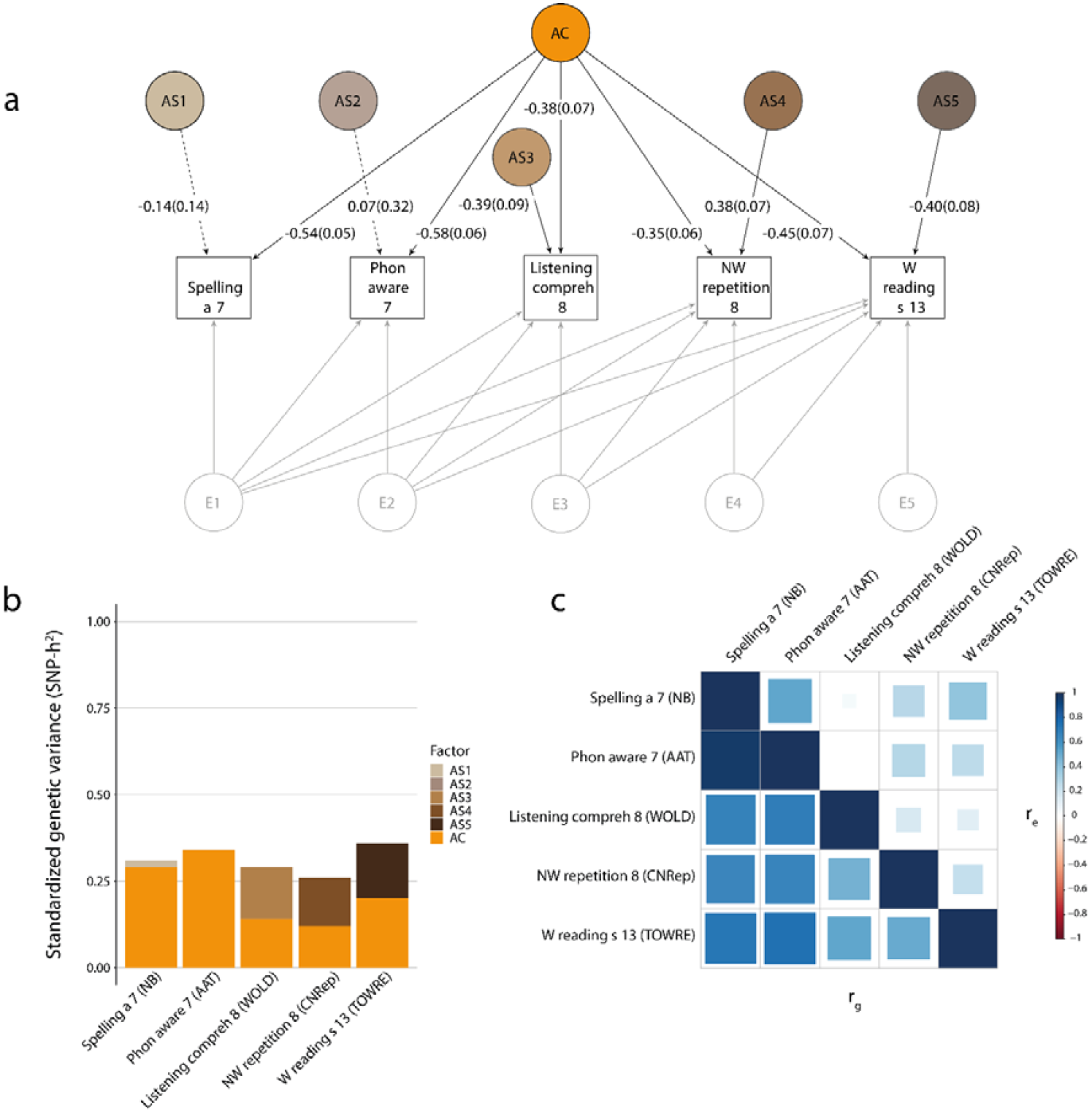
Structural model of literacy-, language-, and PWM-related abilities (Data set: word reading) Genetic-relationship matrix structural equation modeling (GSEM) of five literacy-, language-, and PWM-related abilities) (N=6,383). The path diagram **(a)** depicts the genetic factors of the best-fitting model (IPC model) describing variation in spelling accuracy at age 7 (Spelling a 7, NB), phonemic awareness at age 7 (Phon aware 7, AAT), listening comprehension at age 8 (Listening compreh 8, WOLD), non-word repetition at age 8 (NW repetition 8, CNRep) and word reading speed at age 13 (W reading s 13, TOWRE). The phenotypic variance was dissected into latent common (AC) and specific (AS1-AS5) genetic factors, according to an Independent Pathway model, as well as latent (E1-E5) residual factors, based on a Cholesky decomposition (not shown, Supplementary Table 17, Supplementary Figure 4). Observed measures are represented by squares and latent factors by circles. Single headed arrows (paths) define relationships between variables. Dotted and solid paths represent factor loadings with *p*>0.05 and *p*≤0.05 respectively. The variance of latent variables is constrained to unit variance; this is omitted from the diagram to improve clarity. The variance plot **(b)** depicts the standardized genetic variance components for the model in (a). The correlogram **(c)** shows genetic (r_g_, lower triangle) and residual (r_e_, upper triangle) correlations for the model in (a). Point estimates and their SEs are reported in Supplementary Table 20.

For the passage reading dataset, the IPC model fitted the data best, based on AIC, BIC and LRTs (Table 2, Supplementary Table 13-16). There was evidence for a single shared genetic factor across all literacy-, language-and PWM-related abilities (Figure 2a, Supplementary Table 13), capturing between 13%(SE=0.04) to 45%(SE=0.07) of the phenotypic variance of each trait and, thus, a large proportion of SNP-h^2^ (factorial co-heritabilities > 44%; Supplementary Table 14)(Figure 2b). This is reflected in moderate to strong genetic correlations between traits (r_g_ = 0.49 to 0.97), even between passage reading accuracy and listening comprehension (r_g_=0.66 with SE=0.11, Figure 2c, Supplementary Table 15). The phenotypic variance explained by the latent genetic factor was 27%(SE=0.05) for spelling accuracy, 31%(SE=0.06) for phonemic awareness and 45%(SE=0.07) for passage reading, but only 13%(SE=0.04) for listening comprehension and 14%(SE=0.04) for non-word repetition (Figure 2b, Supplementary Table 13). This corresponds to 91%(SE=0.08), 98%(SE=0.1), 97%(SE=0.08), 44%(SE=0.14), 53%(SE=0.14) of the genetic variance for each trait respectively (Supplementary Table 14). Thus, the majority of genetic variance for literacy-related abilities (>90%) is captured by a shared latent genetic factor. Additional trait-specific factors, capturing here domain-specific influences, were estimated for listening comprehension and non-word repetition (Figure 2a). These factors explained 17%(SE=0.06) and 12%(SE=0.05) of the phenotypic variance respectively (Supplementary Table 13, Figure 2b), corresponding to 56%(SE=0.14) and 47%(SE=0.14) of the respective SNP-h^2^ (Supplementary Table 14).

Identified multivariate model structures for the second dataset including the reading fluency proxy word reading speed (age 13) (Figure 3), assessed approximately four years later than passage reading accuracy (age 9), confirmed the findings above. Model fitting indices suggested a marginally better fit of the IPC model, based on AIC, BIC and LRTs (Table 2, Supplementary Table 17-20). The factor structures identified by the IPC model were consistent with those identified for the first subset and implicated a single genetic factor shared across all domains Figure 3a, (Supplementary Table 17). This latent genetic factor accounted for the majority of genetic variation in several literacy-related skills, such as spelling (94% with SE=0.12) and phonemic awareness (99% with SE=0.13), and approximately half of the SNP-h^2^ in listening comprehension (49% with SE=0.17) and non-word repetition (45% with SE=0.13)(Supplementary Table 18). In contrast to the model for passage reading dataset, the shared factor only captured 56%(SE=0.14) of the SNP-h^2^ for word reading speed (age 13), while the remaining 44%(SE=0.14) were captured by a trait-specific factor (Supplementary Table 18). This pattern mirrors the large specific factor for word reading speed estimated with the Stage 1 reading fluency model (Figure 1a).

Nonetheless, estimated genetic correlations between word reading speed and other phenotypes were still moderate to high; they were 0.52(SE=0.11) with listening comprehension, 0.50(SE=0.10) with non-word repetition, 0.72(SE=0.10) with spelling and 0.74(SE=0.10) with phonemic awareness (Figure 3c). Thus, consistent genetic factor structures between literacy-, language-, and PWM-related abilities (Supplementary Tables 15 and 19) could be robustly identified using different multivariate structural models, irrespective of the genetic composition of the selected reading fluency measure.

As a sensitivity analysis, we confirmed that bivariate genetic correlations between phenotypes, as estimated with GSEM and GCTA, were fully consistent with each other (Supplementary Table 3 and 4). Furthermore, all multivariate GSEM-SNP-h^2^ (Supplementary Table 8, 12, 15, 19) were consistent with univariate GCTA-SNP-h^2^ estimates (Supplementary Table 1), based on derived 95% confidence intervals (CIs).

Lastly, we compared multivariate genetic and residual correlation patterns as fitted by IPC models. Among reading fluency measures (Figure 1c), and, more broadly, literacy-related measures, including spelling accuracy and phonemic awareness (Figure 2c, Figure 3c), strong positive genetic correlations (>0.7) matched modest to strong residual correlations (0.27 to 0.80) with similar directions of effects. Residual correlations among literacy-related measures, with age differences spanning up to six years, were still moderate. For example, residual correlations were estimated at 0.67(SE=0.03) and 0.39(SE=0.06) between spelling accuracy (age 7) and, both passage reading accuracy (age 9)(Figure 2c, Supplementary Table 16) and word reading speed (age 13)(Figure 3c, Supplementary Table 20) respectively. Genetic and residual correlations patterns across different domains, especially oral language and literacy, were more diverse. While there was evidence for moderate to strong genetic correlations across all domains, the respective residual correlations were weak or null (zero within the 95% CIs), as shown in Supplementary Table 16 and 20. Especially, the residual correlations between listening comprehension and literacy-related measures, such as spelling accuracy, phonemic awareness and passage reading, were low in comparison to genetic correlations (Figure 2c); respective residual correlation estimates were 0.08(SE=0.05), 0.02(SE=0.05) and 0.11(SE=0.07), while corresponding genetic correlation estimates were 0.64(SE=0.11), 0.66(SE=0.11) and 0.66 (SE=0.11)(Figure 2c, Supplementary Table 16). The bivariate SNP-h^2^ estimates between these measures were consistent with one, based on derived 95% CIs (Supplementary Table 15).

## Discussion

By modeling multivariate genetic variance within a population-representative sample of unrelated youth using genome-wide markers, we demonstrated that language-, literacy- and working memory-related abilities share a large proportion of their underlying genetic variation, but also reveal substantial differences in their genetic variance composition. These findings imply a pleiotropic set of common genetic variants that contributes broadly to language, literacy and PWM performance and can be augmented by domain-specific genetic variation. Together with evidence for discordant genetic and residual covariance patterns, especially between genetically highly correlated oral language and literacy-related abilities, our findings suggest differences in underlying etiological mechanisms.

### Identification of a shared genetic core factor

In line with long-established findings from multivariate twin research [13-15], we identified an overarching latent genetic (core) factor that is not only shared between word and passage reading abilities, but also with non-reading and other literacy-related abilities that we investigated in this study, including spelling, phonological phonemic awareness, listening comprehension and non-word repetition. Utilizing passage reading accuracy (age 9) as a reading fluency proxy, the overarching genetic factor accounted almost fully for the genetic variance in literacy-related measures (>90%; reading fluency, spelling, phonemic awareness) and, to a lesser extent, for the genetic variance in listening comprehension (44%) and non-word repetition (53%). The identified shared genetic factor suggests wide-spread pleiotropy [30], in support of “generalist genes”, that contribute to multiple shared aspects of cognitive functioning [31]. This includes genetic overlap between listening comprehension and reading fluency, consistent with findings from previous twin research [32, 33], which may arise due to several processes: Not all words can be acquired through phonological decoding, as predicted by the “Simple View of Reading” [6, 7]; words with inconsistent orthographic – phonological mappings need to be memorized or used with contextual cues [33]. Furthermore, once decoding skills have been established and well-practiced on reading words, the orthographic representations of those words become integrated “lexical representations” directly connected to phonological and semantic knowledge [33, 34]. Likewise, phonological awareness is no longer a predictor of reading ability, but instead reciprocally shaped by reading experience [1, 35]. These processes potentially strengthen interrelations between language-and literacy-related abilities with reading experience, as children accumulate detailed orthographic knowledge [1]. In addition, we demonstrated that non-word repetition, considered a proxy of PWM, shares genetic variation with both oral language and literacy-related abilities.

The genetic core factor across multiple literacy-, language-, and PWM-related abilities could be robustly identified using different reading fluency proxies ascertained at different developmental stages, spanning mid-childhood to early adolescence. Based on observational research, word decoding skills have been found to be age-dependent, shaping the transition of “learning to read” to “reading to learn” [22, 36, 37]. In contrast, evidence from analyses of twin samples has demonstrated a high level of genetic stability for word-level decoding [10], with averaged age-to-age (7-12-16 years) genetic correlations of 0.69, consistent with the structural models in this study. Thus, our findings confirm a high level of developmental stability in the genetic architecture of reading fluency using very different methods.

Our analyses also support recent investigations in unrelated individuals, based on genome-wide genotyping information, suggesting that both verbal numerical reasoning and memory-related tasks are related to the same underlying common genetic factor [38], which has long been hypothesized by observational studies [39].

### Identification of trait-specific genetic influences

Beyond a shared genetic core factor, we also observed evidence of at least three instances, where genetic influences capture trait-specific effects that are characteristic of a domain. First, there was a trait-specific genetic variance contribution to oral language abilities (here listening comprehension), consistent with the “Simple View of Reading” [6, 7] distinguishing language comprehension from decoding abilities. This provides converging evidence with recent twin models suggesting a strong patterning of oral language with reading comprehension compared to a distinct and only moderate overlap between oral language with reading fluency [13]. Furthermore, word recognition (decoding) and listening comprehension have been shown to exert independent genetic influences on reading comprehension [40], and children with high reading accuracy and fluency are not necessarily efficient in reading comprehension [41, 42].

Second, we identified a trait-specific genetic variance contribution underlying non-word repetition. This PWM-related trait is thought to consist of two components; (i) a short term phonological store as the non-word has to remain in memory long enough to identify the individual phonemes and their sequence, and (ii) an articulatory rehearsal process as the non-word has to be rehearsed sufficiently rapidly and repeatedly to prevent it from decaying over time [10]. Thus, while these processes are strongly interlinked with language and literacy, supported by findings of this study and others [38], we also find evidence for an independent genetic contribution.

Third, the structural model for literacy-, language-, and PWM-related abilities identified a trait-specific genetic factor for reading fluency as proxied by word reading speed (Figure 3), but not passage reading accuracy (Figure 2). To the best of our knowledge, there currently are no studies that have directly examined the genetic links between passage and word reading fluency. Here, we find evidence that genetic variance in word reading speed can only partially be explained by a common genetic factor shared with other reading fluency measures (27%: Figure 1 and Supplementary Table 6, Stage 1 analysis), and the language-, literacy- and PWM-related core factor (21%: Figure 3 and Supplementary Table 15, Stage 2 analysis). In contrast, variation in passage reading accuracy can almost fully be accounted for by a common genetic factor shared with other reading fluency traits (47%: Figure 1 and Supplementary Table 6, Stage 1 analysis), and the language-, literacy- and PWM-related genetic core factor (45%: Figure 2, Supplementary Table 12, Stage 2 analysis). The differences in genetic structures may arise as reading accuracy and speed measures, based on grammatically and semantically coherent passages, are likely to be more influenced by comprehension skills compared to those based on single-word reading [1, 43].

Given the near-fully capture of genetic variance underlying literacy-related abilities (>90%) such as passage reading fluency, spelling and phonemic awareness by the genetic core factor, it is possible that domain-specific genetic factors cover a range of cognitive abilities with genetic mechanisms that are unique to oral language, PWM and word decoding abilities, consistent with interrelated, but different genetic etiologies.

### Discordant genetic and residual correlation patterns

Irrespective of strong genetic relationships between language and literacy-related traits, the corresponding residual correlations were approximately zero. This lack of residual overlap with oral language was observed for literacy-related measures ascertained at different ALSPAC clinics, both before and after the assessment of oral language. In contrast, literacy-related measures were, without exception, weakly to strongly residually intercorrelated, using different psychological instruments and raters across a window of approximately six years, rendering correlated assessment-specific error unlikely. Furthermore, the polygenic nature of both oral language and literacy skills, supported by large SNP-h^2^ estimates, suggests little impact of rare genetic influences, due to the low power of population-based cohorts to detect rare genetic effects [44]. Thus, our results are consistent with differences in underlying etiological mechanisms that may manifest through their (non-genetic) environmental interrelationships. Our findings suggest reduced modifiability of oral language through literacy, potentially affecting the scope of centrally applied reading intervention programs, and add further support for the transferability of acquired skills within the literacy domain. Latter is strongly supported by twin study findings of bi-directional effects between reading fluency and reading comprehension, and overlapping genetic and environmental factors [45].

### Strengths and limitations

This study benefits from the modeling of multivariate genetic variances in unrelated individuals based on directly assessed genome-wide genotyping information and a likelihood-based comparison of multiple model structures against saturated (and computationally expensive) Cholesky models. As such we combine here twin research-based structural equation modeling approaches popular in multivariate twin studies with genomic approaches using GRMs. A major strength is introducing novel IPC models that jointly enhance both the interpretability of model coefficients and the accuracy of estimates, while incorporating information from all available data. Furthermore, data from ALSPAC has a large sample size for a comprehensive set of literacy-, language-, and PWM-related measures.

Our study has several limitations: firstly, literacy-, language- and PWM-related abilities were assessed during mid-childhood and adolescence, ALSPAC lacked longitudinal information to determine whether there are development-specific changes in the genetic architecture contributing to these traits. Population-level phenomena, such as assortative mating and dynastic effects [46, 47] can potentially inflate genetic correlations between assessed measures and upward-bias SNP heritability, even in seemingly unrelated individuals [47]. For example, parents may select each other based on their compatibility in cognitive skills which may create a reading-rich environment for their children. These phenomena could explain some shared genetic variation across measures in our study though they are unlikely to account for the entirety of shared genetic variance. Population-level biases should systematically affect all intercorrelated traits and are, thus, unlikely to account for differences in genetic and residual correlation patterns between language and literacy measures. Furthermore, the large variation in genetic factor loadings across language-, literacy- and PWM-domain suggests differences in underlying etiological mechanisms. Finally, the lack of independent cohorts with comparable sets of literacy-, language- and PWM-related measures prevents a direct replication of our work, although the consistency of our findings with many recent twin study reports supports the validity of our results.

## Conclusions

Modeling multivariate genetic architectures as captured by genome-wide variation, we have identified a core genetic factor that is shared among measures of literacy, language and PWM, and augmented by trait-specific genetic variance contributions, especially for oral language and PWM. Together with evidence for discordant genetic and residual interrelationships between oral language and literacy, our findings suggest a diverse spectrum of interrelated cognitive skills involving multiple etiological mechanisms and different levels of trait modifiability.

## Methods

### Study participants

#### Cohort description

This study was carried out using longitudinal data from children and adolescents between 7 and 13 years from the ALSPAC study, a UK population-based pregnancy-ascertained birth-cohort (estimated birth date: 1991-1992)[48, 49]. Informed consent for the use of data collected via questionnaires and clinics was obtained from participants following the recommendations of the ALSPAC Ethics and Law Committee at the time. Consent for biological samples has been collected in accordance with the Human Tissue Act (2004). Details on the recruitment of the ALSPAC cohort can be found in Supplementary Note 1.

Please note that the study website contains details of all the data that is available through a fully searchable data dictionary and variable search tool: http://www.bristol.ac.uk/alspac/researchers/our-data/.

### Measures of literacy-, language-, and PWM-related abilities

#### Phenotype descriptions

The literacy-, language-, and PWM-related abilities in this study included word and passage reading abilities (accuracy and speed), reading-related decoding abilities (non-word reading: accuracy and speed), spelling abilities (accuracy), phonemic awareness, listening comprehension, and non-word repetition as assessed with both standardized and ALSPAC-specific instruments (Table 1). We excluded ALSPAC measures which were composites of multiple language and literacy domains (such as e.g. verbal intelligence) or which involved tiered assessments.

##### Reading accuracy age 9 (NBO), words

To evaluate word reading accuracy, the child was asked to read aloud a list of 10 real words. Nunes and Bryant selected these real words, which are a subset of those proposed by Nunes, Bryant and Olsen (NBO)[50]. The reading accuracy score consists of the total number of items that the child read correctly. The test-retest reliability of word reading was 0.80, and had a correlation of 0.85 with the Schonell Word Reading Task[51].

##### Reading speed and accuracy age 9 (NARA II), passages

Children’s passage reading speed and accuracy were assessed with the revised Neale Analysis of Reading Ability[52] (NARA II). A story was given to the child to read. The tester recorded the time it took the child to read each passage, and noted any errors made by the child. All scores were standardized by age.

##### Reading speed age 13 (TOWRE), words

The Test of Word Reading Efficiency[53] (TOWRE) was used to assess overall word reading efficiency using the Sight Word Efficiency sub-scale. The child was given 45 seconds to read as many words as possible. Words that were skipped, or wrong words were marked by the tester. The reading speed score was computed as the total number of correct words read by the child.

##### Non-word reading accuracy age 9 (NBO)

The child was asked to read aloud 10 non-words. These words had been selected from a larger selection of non-words on a ALSPAC-specific instrument based on the research conducted by Nunes and colleagues[50]. The test - retest reliability of the non-word reading task was 0.73. The tester emphasized to the child that the words ware made-up and asked the child to read all the non-words in the way that they thought they should be read. A total score was computed as the sum of the number of items read correctly by the child based on regular symbol-sound correspondences of written English.

##### Non-word reading speed age 13 (TOWRE)

The Test of Word Reading Efficiency [53] (TOWRE) was also used to assess non-word reading speed using the Phonemic Decoding Efficiency sub-scale. It is designed as a relatively pure measure of decoding, independent of meaning. The child had 45 seconds to read as many non-words as possible. Non-words that a child skipped or got wrong were marked by the tester. A total score was computed as the sum of the number of correct non-words read by the child based on regular symbol-sound correspondences of written English.

##### Spelling accuracy age 7 and age 9 (NB)

Spelling accuracy was assessed by asking a child to spell a series of 15 words. The words were chosen specifically for this age group after piloting on several hundred children (Nunes and Bryant, ALSPAC-specific measure [15]). The words included regular and irregular words of differing frequencies and were put in an order of increasing difficulty. For each word, the tester first read the word out alone to the child, then within a specific sentence incorporating the word, and finally alone again. The child was then asked to write down the spelling. The spelling accuracy score is computed as the number of words spelt correctly.

##### Phonemic awareness age 7 (AAT)

Phonemic awareness was assessed using the Auditory Analysis Test [54] (AAT). The task contained two practice and 40 test items of increasing difficulty. For each item, the child was first asked to repeat the word, and then to produce it again but with part of the word (a phoneme or several phonemes) removed. For example, the word to repeat initially is “sour” and then the child is asked to repeat it again without /s/ to which the correct response is “our”. The test assessed seven omission categories: the omission of a first, medial or final syllable, the omission of the initial consonant, the omission of the final consonant of a one-syllable word, and the omission of the first consonant or consonant blend of a medial consonant. The words from different categories were mixed. A total score was computed as the sum of correct responses over all types of omission.

##### Listening comprehension at 8 (WOLD)

Listening comprehension is a subtest of the Wechsler Objective Language Dimensions (WOLD) [55]. A picture was shown to the participant, and the examiner read aloud a paragraph about the picture. The participant then answered multiple open-ended questions on what they have heard.

##### Non-word repetition at age 8 (NWR)

Children’s short-term memory was evaluated by a non-word repetition test [11]. The test contains 12 nonsense words, for each of 3, 4 and 5 syllables, all conforming to English rules for sound combinations. The participant was asked to listen to each of these 36 words from an audio cassette recorder and then repeat each word.

#### Phenotype transformations

Any children who stopped prematurely during the psychological tests were excluded from the study. The scores of each test were adjusted for age, sex and the two principal components from the genotyping analysis to correct for population stratification [56]. The scores from passage reading measures (NARA) are already age-standardized, and therefore these scores were not further adjusted for age. All residualized scores were finally rank-transformed to improve the GSEM log-likelihood estimation assuming multivariate normality.

SNP-h^2^ estimates (Supplementary Table 1) and bivariate phenotypic correlations (based on Pearson correlation coefficients, Supplementary Table 2) between measures, using both untransformed and rank-transformed scores, were comparable with each other and with previous reports [15].

### Genotyping and imputation

ALSPAC children were genotyped using Illumina HumanHap550 quad-chip platform. The ALSPAC GWAS data were generated by Sample Logistics and Genotyping Facilities at the Wellcome Trust Sanger Institute and LabCorp (Laboratory Corporation of America) using support from 23andMe. After quality control (individual call rate>0.97, SNP call rate>0.95, Minor allele frequency (MAF)>0.01, Hardy-Weinberg equilibrium (HWE) *p*>10^−7^, and removal of individuals with cryptic relatedness and non-European ancestry), genome-wide data were available for 8,237 children and 465,740 directly genotyped Single Nucleotide Polymorphisms (SNPs).

### Genetic-relationship matrix structural equation modeling

The multivariate genetic architecture of literacy-, language-, and PWM-related abilities was modelled using genetic-relationship matrix structural equation modeling (R:gsem library, version 1.1.0, https://gitlab.gwdg.de/beate.stpourcain/gsem) [27]. This multivariate analysis technique combines whole-genome genotyping information with structural equation modeling techniques applied in twin research [57] to model multivariate genetic variances, using a maximum likelihood approach [27]. Thus, within the GSEM framework genetic and environmental influences are modelled as latent factors. Specifically, phenotypic variance for each measure was dissected into genetic and residual influences (AE model) using a full Cholesky decomposition and an independent pathway model [27, 58]:

i. The Cholesky decomposition model is a fully parametrised descriptive model without any restrictions on the structure of latent genetic and residual influences. This saturated model can be fit to the data through decomposition of both the genetic variance and residual variance into as many latent factors as there are observed variables.
ii. The independent pathway model specifies a common genetic and a common residual factor, in addition to trait-specific genetic and residual influences.
iii. The combined Cholesky decomposition and independent pathway (IPC) model structures the genetic variance as an independent pathway model (consisting of common and measurement specific influences) and the residual variance as a Cholesky decomposition model (where the number of latent factors are the same as the number of observed variables). Supplementary Figure 3 shows an example of an IPC model with six traits, and Supplementary Figure 4 an example of an IPC model with five traits.

The goodness-of-fit of GSEM models to the data was evaluated with LRTs, AIC [59] and BIC [60].

In the present study, we modelled rank-transformed residualized phenotypes (adjusted for age, sex and the first two principal components) as described above. Genetic-relationship matrices were constructed based on directly genotyped variants in unrelated individuals, using GCTA software [28]. Individuals with a relatedness > 0.05 (off-diagonals within a genetic-relatedness matrix) were excluded. Factor loadings were evaluated using Wald tests.

Multivariate models in unrelated individuals, studying interindividual genetic variation captured by genome-wide genetic variation, are computationally expensive [27]. For example, a 6-factor Cholesky decomposition model, as fitted within this study, can require 6 weeks computing time even on a system incorporating at least 4 parallel cores of 3 GHz and between 50 to 75 Gb memory. Hence, we started the modeling process by strategically combining similar measures, such as reading fluency or spelling abilities, to reduce the number of studied instruments by selecting proxies that most comprehensively capturing the shared genetic variance of an entire domain. The advantage of this approach is that we can control the extent to which selected proxies represent underlying shared genetic factors, either fully or along with specific measurement-related influences, and subsequently assess the stability of the identified genetic structures. The identified proxy measures in Stage 1 were eventually jointly modelled with other measures as part of Stage 2 analysis, with models across three phenotypic domains: literacy-related measures (reading fluency, spelling, phonemic awareness), oral language (listening comprehension) and PWM (non-word repetition).

Applying a conservative approach, we also evaluated all derived factor loadings of the Stage 2 model against an experiment-wide error rate of 0.007, estimated based on the effective number of individually analyzed phenotypes using Matrix Spectral Decomposition (MatSpD) [61], accounting for all previously conducted statistical assessments. However, a correction for multiple testing is not directly applicable, as we jointly analyze language-and literacy related traits with different genetic architectures using multivariate modeling with GSEM.

Identified structural models were used to estimate SNP-h^2^ as well as genetic correlations, factorial co-heritabilities (the proportion of total trait genetic variance explained by a specific genetic factor) and bivariate heritabilities (the contribution of genetic covariance to the observed phenotypic covariance between two measures), as defined here:

Bivariate genetic correlation between phenotypes, measuring the extent to which two phenotypes 1 and 2 share genetic factors (ranging from −1 to 1), can be derived using estimated genetic variances and covariances [62] according to:

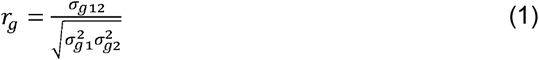

where σ_g12_ is the genetic covariance between phenotypes 1 and 2, and σ^2^ _g1_ and σ^2^ _g2_ are the respective genetic variances.

A measure of factorial co-heritability was introduced to assess the relative contribution of a genetic factor to the genetic variance of a phenotype, estimated with the gsem package (R:gsem library, version 1.1.0). The factorial co-heritability f_g_^2^ is defined as:

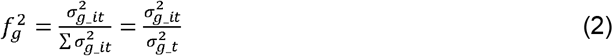

where σ^2^_g_it_ is the genetic variance of the genetic factor i contributing to trait t and σ^2^ _g_t_ the total genetic variance of trait t, based on standardised factor loadings. Corresponding SEs were derived using the Delta method.

Bivariate heritability [63] was defined as the ratio of the genetic to the phenotypic covariance between two traits, and was estimated using the gsem package (R:gsem library, version 1.1.0).

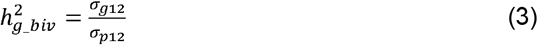

where *σ*_*g*12_ is the genetic covariance, estimated based on unstandardized path coefficients, and *σ*_*p*12_ the phenotypic covariance from observed rank-transformed measures. The respective SEs were approximated by the SE of the genetic covariance divided by the phenotypic covariance (as the SE of the phenotypic covariance is very small).

### Genetic-relationship-matrix residual maximum likelihood

The GCTA software package (v1.25.2) [28] can be used to estimate the proportion of phenotypic variation that is explained by markers on genotyping chip arrays using genetic-relationship-matrix residual maximum likelihood (GREML) [28] (AE model). Likewise, bivariate GREML [64] can be applied to estimate genetic covariances and genetic correlations between two phenotypes.

Univariate and bivariate GREML were carried out as part of sensitivity analyses.

## Data availability

The data used are available through a fully searchable data dictionary (http://www.bristol.ac.uk/alspac/researchers/our-data/). Access to ALSPAC data can be obtained as described within the ALSPAC data access policy (http://www.bristol.ac.uk/alspac/researchers/access/). All analyses were performed using freely accessible software. Requests for scripts or other analysis details can be sent via email to the corresponding authors.

## Acknowledgements

We are extremely grateful to all the families who took part in this study, the midwives for their help in recruiting them, and the whole ALSPAC team, which includes interviewers, computer and laboratory technicians, clerical workers, research scientists, volunteers, managers, receptionists and nurses. The UK Medical Research Council and Wellcome (Grant ref: 217065/Z/19/Z) and the University of Bristol provide core support for ALSPAC. GWAS data was generated by Sample Logistics and Genotyping Facilities at Wellcome Sanger Institute and LabCorp (Laboratory Corporation of America) using support from 23andMe. A comprehensive list of grants funding is available on the ALSPAC website (http://www.bristol.ac.uk/alspac/external/documents/grant-acknowledgements.pdf). CYS works in a unit that receives support from the University of Bristol and the UK Medical Research Council. EV, BSTP and SEF are supported by the Max Planck Society. BSTP is also supported by the Simons Foundation (Award ID: 514787). This publication is the work of the authors and CYS will serve as guarantors for the contents of this paper.

## Author contributions

BSTP developed the study concept and, CYS and EV contributed to the study design. CYS and BSTP performed the data analysis and CYS carried out the interpretation under the supervision of BSTP. CYS and BSTP drafted the manuscript and, EV, GDS, SEF, BV and PSD provided critical revisions. All authors approved the final version of the manuscript for submission.

