## Supplementary Material for "The Multivariate Genome-wide Architecture of Interrelated Literacy, Language and Working Memory Skills Reveals Distinct Etiologies"

### Supplementary Notes

#### Supplementary Note 1: ALSPAC cohort description

Pregnant women resident in Avon, UK with expected dates of delivery 1st April 1991 to 31st December 1992 were invited to take part in the study. The initial number of pregnancies enrolled is 14,541 (for these at least one questionnaire has been returned or a “Children in Focus” clinic had been attended by 19/07/99). Of these initial pregnancies, there was a total of 14,676 fetuses, resulting in 14,062 live births and 13,988 children who were alive at 1 year of age.

When the oldest children were approximately 7 years of age, an attempt was made to bolster the initial sample with eligible cases who had failed to join the study originally. As a result, when considering variables collected from the age of seven onwards (and potentially abstracted from obstetric notes) there are data available for more than the 14,541 pregnancies mentioned above. The number of new pregnancies not in the initial sample (known as Phase I enrolment) that are currently represented on the built files and reflecting enrolment status at the age of 24 is 913 (456, 262 and 195 recruited during Phases II, III and IV respectively), resulting in an additional 913 children being enrolled. The phases of enrolment are described in more detail in the cohort profile paper and its update [1, 2]. The total sample size for analyses using any data collected after the age of seven is therefore 15,454 pregnancies, resulting in 15,589 fetuses. Of these 14,901 were alive at 1 year of age.

A 10% sample of the ALSPAC cohort, known as the Children in Focus (CiF) group, attended clinics at the University of Bristol at various time intervals between 4 to 61 months of age. The CiF group were chosen at random from the last 6 months of ALSPAC births (1432 families attended at least one clinic). Excluded were those mothers who had moved out of the area or were lost to follow-up, and those partaking in another study of infant development in Avon.

Please note that the study website contains details of all the data that is available through a fully searchable data dictionary and variable search tool: <http://www.bristol.ac.uk/alspac/researchers/our-data/>

### Supplementary Tables

| **Measure** | **N** | **GCTA SNP-h^2^ (SE)** | |
| --- | --- | --- | --- |
|  |  | **Residuals** | **Rank-transformed**  **residuals** |
| P reading a 9 (NARA) | 5048 | 0.5029 (0.07) | 0.5029 (0.07) |
| P reading s 9 (NARA) | 5037 | 0.4453 (0.07) | 0.4453 (0.07) |
| W reading a 9 (NBO) | 5574 | 0.3620 (0.06) | 0.4555 (0.06) |
| W reading s 13 (TOWRE) | 4131 | 0.4001 (0.09) | 0.4024 (0.09) |
| NW reading a 9 (NBO) | 5569 | 0.2954 (0.06) | 0.3213 (0.06) |
| NW reading s 13 (TOWRE) | 4121 | 0.3512 (0.09) | 0.3774 (0.09) |
| Spelling a 7 (NB) | 5637 | 0.3286 (0.06) | 0.3170 (0.06) |
| Spelling a 9 (NB) | 5564 | 0.3633 (0.07) | 0.3786 (0.06) |
| Phon aware 7 (AAT) | 5749 | 0.3861 (0.06) | 0.3852 (0.06) |
| Listening compreh 8 (WOLD) | 5324 | 0.3180 (0.07) | 0.3182 (0.07) |
| NW repetition 8 (CNRep) | 5315 | 0.3294 (0.07) | 0.3168 (0.07) |

#### Supplementary Table 1: SNP-heritability estimates for residual and rank transformed literacy‑, language-, and PWM-related abilities

SNP-heritability (SNP-h^2^) was estimated with Genetic-relationship-matrix Restricted Maximum Likelihood analysis (GREML) using Genome-wide Complex Trait Analysis (GCTA) software. Residuals, residuals of language or literacy-related measures after adjustment for sex, age and the top two most significant principal components, unless the measure was age normalised. Rank-transformed residuals, rank-transformed residuals of language or literacy-related measures. PWM, Phonological working memory

#### Supplementary Table 2: Phenotypic correlations between literacy-, language-, and PWM-related abilities

|  | P reading a 9 | P reading s 9 | W reading a 9 | W reading s 13 | NW reading a 9 | NW reading s 13 | Spelling a 7 | Spelling a 9 | Phon aware 7 | Listening compreh 8 | NW repetition 8 |
| --- | --- | --- | --- | --- | --- | --- | --- | --- | --- | --- | --- |
| P reading a 9 (NARA II) | 1 | 0.74 | 0.76 | 0.62 | 0.56 | 0.65 | 0.63 | 0.65 | 0.48 | 0.3 | 0.34 |
| P reading s 9 (NARA II) | 0.74 | 1 | 0.63 | 0.5 | 0.72 | 0.56 | 0.67 | 0.71 | 0.57 | 0.23 | 0.36 |
| W reading a 9 (NBO) | 0.76 | 0.63 | 1 | 0.59 | 0.70 | 0.65 | 0.75 | 0.78 | 0.64 | 0.29 | 0.44 |
| W reading s 13 (TOWRE) | 0.62 | 0.5 | 0.60 | 1 | 0.45 | 0.81 | 0.48 | 0.52 | 0.41 | 0.24 | 0.29 |
| NW reading a 9 (NBO) | 0.56 | 0.72 | 0.70 | 0.45 | 1 | 0.53 | 0.63 | 0.66 | 0.55 | 0.17 | 0.33 |
| NW reading s 13 (TOWRE) | 0.65 | 0.57 | 0.65 | 0.81 | 0.53 | 1 | 0.54 | 0.58 | 0.45 | 0.20 | 0.28 |
| Spelling a 7 (NB) | 0.62 | 0.66 | 0.74 | 0.47 | 0.62 | 0.53 | 1 | 0.77 | 0.66 | 0.19 | 0.35 |
| Spelling a 9 (NB) | 0.65 | 0.71 | 0.78 | 0.51 | 0.66 | 0.58 | 0.75 | 1 | 0.59 | 0.21 | 0.36 |
| Phon aware 7 (AAT) | 0.48 | 0.57 | 0.64 | 0.40 | 0.54 | 0.45 | 0.66 | 0.58 | 1 | 0.19 | 0.38 |
| Listening compreh 8 (WOLD) | 0.31 | 0.23 | 0.29 | 0.24 | 0.17 | 0.20 | 0.20 | 0.20 | 0.19 | 1 | 0.23 |
| NW repetition 8 (CNRep) | 0.34 | 0.36 | 0.43 | 0.29 | 0.32 | 0.28 | 0.35 | 0.36 | 0.38 | 0.23 | 1 |

Phenotypic correlations are Pearson’s correlation coefficients. Correlations between residualized scores (adjusted for sex, age and top two most significant principal components) are shown in the lower triangle and correlations among rank-transformed residualized scores within the upper triangle.

AAT, Auditory Analysis Test; ALSPAC, Avon Longitudinal study of Parents and Children; CNRep, Children’s Test of Nonword Repetition; NARA II, The Neale Analysis of Reading Ability-Second Revised British Edition; NBO, ALSPAC-specific assessment developed by Nunes, Bryant and Olson; NB, ALSPAC-specific assessment developed by Nunes and Bryant; TOWRE, Test Of Word Reading Efficiency; PWM, Phonological working memory; WOLD, Wechsler Objective Language Dimensions; WORD, Wechsler Objective Reading Dimension

#### Supplementary Table 3: Genetic correlations between literacy-, language-, and PWM-related abilities (GCTA)

|  | P reading a 9 | P reading s 9 | W reading a 9 | W reading s 13 | NW reading a 9 | NW reading s 13 | Spelling a 7 | Spelling a 9 | Phon aware 7 | Listening compreh 8 | NW repetition 8 |
| --- | --- | --- | --- | --- | --- | --- | --- | --- | --- | --- | --- |
| P reading a 9 (NARA II) | 1 |  |  |  |  |  |  |  |  |  |  |
| P reading s 9 (NARA II) | 0.93 (0.04) | 1 |  |  |  |  |  |  |  |  |  |
| W reading a 9 (NBO) | 0.89 (0.04) | 0.84 (0.06) | 1 |  |  |  |  |  |  |  |  |
| W reading s 13 (TOWRE) | 0.90 (0.08) | 0.94 (0.08) | 0.67 (0.09) | 1 |  |  |  |  |  |  |  |
| NW reading a 9 (NBO) | 0.90 (0.05) | 0.81 (0.08) | 0.98 (0.05) | 0.68 (0.11) | 1 |  |  |  |  |  |  |
| NW reading s 13 (TOWRE) | 0.93 (0.07) | 0.92 (0.07) | 0.92 (0.08) | 0.87 (0.04) | 0.96 (0.11) | 1 |  |  |  |  |  |
| Spelling a 7 (NB) | 0.96 (0.05) | 0.82 (0.07) | 0.87 (0.06) | 0.71 (0.11) | 0.92 (0.08) | 0.87 (0.10) | 1 |  |  |  |  |
| Spelling a 9 (NB) | 0.96 (0.03) | 0.90 (0.06) | 0.95 (0.04) | 0.83 (0.09) | 0.92 (0.06) | 0.89 (0.08) | 0.97 (0.04) | 1 |  |  |  |
| Phon aware 7 (AAT) | 0.97 (0.06) | 0.86 (0.09) | 0.86 (0.07) | 0.77 (0.12) | 0.93 (0.08) | 0.86 (0.11) | 0.98 (0.05) | 0.87 (0.07) | 1 |  |  |
| Listening compreh 8 (WOLD) | 0.67 (0.11) | 0.80 (0.12) | 0.49 (0.12) | 0.58 (0.14) | 0.46 (0.14) | 0.49 (0.15) | 0.73 (0.14) | 0.55 (0.13) | 0.64 (0.13) | 1 |  |
| NW repetition 8 (CNRep) | 0.73 (0.10) | 0.69 (0.12) | 0.61 (0.10) | 0.41 (0.14) | 0.72 (0.12) | 0.40 (0.15) | 0.66 (0.12) | 0.81 (0.11) | 0.70 (0.11) | 0.54 (0.14) | 1 |

Bivariate genetic correlations were estimated with Genetic-relationship-matrix Restricted Maximum Likelihood analysis (REML), using Genome-wide complex trait analysis (GCTA) software. Standard errors are in brackets. All analyses are based rank-transformed residualized scores.

AAT, Auditory Analysis Test; ALSPAC, Avon Longitudinal study of Parents and Children; CNRep, Children’s Test of Nonword Repetition; NARA II, The Neale Analysis of Reading Ability-Second Revised British Edition; NBO, ALSPAC-specific assessment developed by Nunes, Bryant and Olson; NB, ALSPAC-specific assessment developed by Nunes and Bryant; TOWRE, Test Of Word Reading Efficiency; PWM, Phonological working memory; WOLD, Wechsler Objective Language Dimensions; WORD, Wechsler Objective Reading Dimension

#### Supplementary Table 4: Genetic correlations between literacy-, language-, and PWM-related abilities (GSEM)

|  | P reading a 9 | P reading s 9 | W reading a 9 | W reading s 13 | NW reading a 9 | NW reading s 13 | Spelling a 7 | Spelling a 9 | Phon aware 7 | Listening compreh 8 | NW repetition 8 |
| --- | --- | --- | --- | --- | --- | --- | --- | --- | --- | --- | --- |
| P reading a 9 (NARA II) | 1 |  |  |  |  |  |  |  |  |  |  |
| P reading s 9 (NARA II) | 0.93  (0.04) | 1 |  |  |  |  |  |  |  |  |  |
| W reading a 9 (NBO) | 0.89 (0.04) | 0.84 (0.06) | 1 |  |  |  |  |  |  |  |  |
| W reading s 13 (TOWRE) | 0.84 (0.07) | 0.89 (0.07) | 0.66  (0.09) | 1 |  |  |  |  |  |  |  |
| NW reading a 9 (NBO) | 0.93 (0.05) | 0.81 (0.08) | 0.98  (0.05) | 0.67  (0.11) | 1 |  |  |  |  |  |  |
| NW reading s 13 (TOWRE) | 0.87 (0.06) | 0.87 (0.07) | 0.90  (0.08) | 0.88  (0.04) | 0.95  (0.11) | 1 |  |  |  |  |  |
| Spelling a 7 (NB) | 0.95 (0.05) | 0.81 (0.08) | 0.87  (0.06) | 0.70  (0.11) | 0.92  (0.08) | 0.85  (0.10) | 1 |  |  |  |  |
| Spelling a 9 (NB) | 0.96 (0.04) | 0.90 (0.06) | 0.95  (0.04) | 0.82  (0.09) | 0.92  (0.06) | 0.87  (0.08) | 0.97  (0.05) | 1 |  |  |  |
| Phon aware 7 (AAT) | 0.97 (0.06) | 0.85 (0.09) | 0.85  (0.07) | 0.76  (0.12) | 0.92  (0.09) | 0.84  (0.11) | 0.98  (0.06) | 0.86  (0.07) | 1 |  |  |
| Listening compreh 8 (WOLD) | 0.66 (0.11) | 0.79 (0.13) | 0.49  (0.12) | 0.57  (0.14) | 0.46  (0.14) | 0.49  (0.15) | 0.72  (0.15) | 0.55  (0.14) | 0.62  (0.13) | 1 |  |
| NW repetition 8 (CNRep) | 0.72 (0.10) | 0.69 (0.13) | 0.61  (0.11) | 0.41  (0.14) | 0.72  (0.13) | 0.40  (0.15) | 0.65  (0.13) | 0.81  (0.12) | 0.69  (0.11) | 0.53  (0.14) | 1 |

Bivariate genetic correlations were estimated with Genetic-relationship matrix structural equation modeling (GSEM). Standard errors are in brackets. All analyses are based rank-transformed residualized scores.

AAT, Auditory Analysis Test; ALSPAC, Avon Longitudinal study of Parents and Children; CNRep, Children’s Test of Nonword Repetition; NARA II, The Neale Analysis of Reading Ability-Second Revised British Edition; NBO, ALSPAC-specific assessment developed by Nunes, Bryant and Olson; NB, ALSPAC-specific assessment developed by Nunes and Bryant; TOWRE, Test Of Word Reading Efficiency; PWM, Phonological working memory; WOLD, Wechsler Objective Language Dimensions; WORD, Wechsler Objective Reading Dimension

#### Supplementary Table 5: Modeling strategy

| **Stage** | **Traits** | **Measures within model** | **Structural model** |
| --- | --- | --- | --- |
| **1** | **Reading fluency** | NW reading a 9 (NBO);  NW reading s 13 (TOWRE);  W reading a 9 (NBO);  W reading s 13 (TOWRE);  P reading a 9 (NARA);  P reading s 9 (NARA) | IPC, Cholesky,  Independent pathway |
| **1** | **Spelling** | Spelling a 7 (NB);  Spelling a 9 (NB) | Cholesky |
| **2** | **Literacy-, language-, and PWM-related abilities**  (Data set: passage reading) | Spelling a 7 (NB);  Phon aware 7 (AAT);  Listening compreh 8 (WOLD);  NW repetition 8 (CNRep);  **P reading a 9 (NARA)** | IPC, Cholesky,  Independent pathway |
| **2** | **Literacy-, language-, and PWM-related abilities**  (Data set: word reading) | Spelling a 7 (NB);  Phon aware 7 (AAT);  Listening compreh 8 (WOLD);  NW repetition 8 (CNRep);  **W reading s 13 (TOWRE)** | IPC, Cholesky,  Independent pathway |

AAT, Auditory Analysis Test; ALSPAC, Avon Longitudinal study of Parents and Children; CNRep, Children’s Test of Nonword Repetition; NARA II, The Neale Analysis of Reading Ability-Second Revised British Edition; NBO, ALSPAC-specific assessment developed by Nunes, Bryant and Olson; NB, ALSPAC-specific assessment developed by Nunes and Bryant; TOWRE, Test Of Word Reading Efficiency; PWM, Phonological working memory; WOLD, Wechsler Objective Language Dimensions; WORD, Wechsler Objective Reading Dimension; IPC, Combined Independent Pathway (genetic part) / Cholesky (residual part) model

#### Supplementary Table 6: GSEM IPC model for reading fluency, standardised factor loadings

| **Label** | **Path coefficient** | | **Variance explained (%)** |
| --- | --- | --- | --- |
|  | **Estimate (SE)** | **p** | **Estimate (SE)** |
| ac1 | -0.55 (0.05) | 2.39 x 10^-24^ | 30.1 (0.06) |
| ac2 | -0.58 (0.06) | 9.00 x 10^-21^ | 33.7 (0.07) |
| ac3 | -0.64 (0.05) | 1.57 x 10^-36^ | 40.8 (0.07) |
| ac4 | -0.51 (0.07) | 2.64 x 10^-13^ | 26.6 (0.07) |
| ac5 | -0.68 (0.05) | 6.17 x 10^-49^ | 46.5 (0.07) |
| ac6 | -0.60 (0.05) | 4.44 x 10^-28^ | 35.5 (0.07) |
| as1 | <0.01 (0.10) | 1.00 | <0.01 (<0.01) |
| as2 | -0.18 (0.08) | 0.02 | 3.1 (0.03) |
| as3 | -0.20 (0.06) | 9.21 x 10^-04^ | 3.8 (0.02) |
| as4 | 0.24 (0.06) | 9.44 x 10^-05^ | 5.8 (0.03) |
| as5 | -0.18 (0.05) | 9.80 x 10^-04^ | 3.2 (0.02) |
| as6 | 0.17 (0.09) | 0.05 | 2.9 (0.03) |
| e11 | -0.84 (0.04) | 6.78 x 10^-123^ | 69.8 (0.06) |
| e21 | -0.27 (0.05) | 2.38 x 10^-07^ | 7.4 (0.03) |
| e31 | -0.44 (0.05) | 1.64 x 10^-18^ | 19.5 (0.04) |
| e41 | -0.23 (0.05) | 6.42 x 10^-06^ | 5.4 (0.02) |
| e51 | -0.39 (0.05) | 5.82 x 10^-16^ | 15.3 (0.04) |
| e61 | -0.27 (0.05) | 6.94 x 10^-09^ | 7.6 (0.03) |
| e22 | -0.75 (0.04) | 8.19 x 10^-97^ | 57.7 (0.05) |
| e32 | -0.13 (0.03) | 5.81 x 10^-05^ | 1.8 (0.01) |
| e42 | -0.62 (0.04) | 8.85 x 10^-47^ | 38.7 (0.05) |
| e52 | -0.22 (0.04) | 3.70 x 10^-08^ | 4.8 (0.02) |
| e62 | -0.29 (0.04) | 1.08 x 10^-11^ | 8.6 (0.03) |
| e33 | -0.58 (0.02) | 7.42 x 10^-129^ | 33.7 (0.03) |
| e43 | -0.04 (0.03) | 0.15 | 0.1 (<0.01) |
| e53 | -0.21 (0.03) | 8.54 x 10^-13^ | 4.2 (0.01) |
| e63 | -0.13 (0.03) | 2.39 x 10^-05^ | 1.8 (0.01) |
| e44 | 0.49 (0.03) | 1.73 x 10^-72^ | 24.7 (0.03) |
| e54 | 0.07 (0.03) | 7.26 x 10^-03^ | 0.5 (<0.01) |
| e64 | 0.14 (0.03) | 1.86 x 10^-06^ | 1.9 (0.01) |
| e55 | -0.50 (0.02) | 1.44 x 10^-121^ | 25.1 (0.02) |
| e65 | -0.24 (0.03) | 7.11 x 10^-18^ | 5.8 (0.01) |
| e66 | -0.60 (0.02) | 7.63 x 10^-128^ | 36.3 (0.03) |

GSEM IPC (best-fitting) model for reading fluency (Figure 1, N=5,866). The model consists of (in that order of traits 1 to 6): NW reading a 9 (NBO), NW reading s 13 (TOWRE), W reading a 9 (NBO), W reading s 13 (TOWRE), P reading a 9 (NARA) and P reading s 9 (NARA). The phenotypic variance was dissected into common (AC1, AC2, AC3, AC4, AC5 and AC6), and specific genetic (AS1, AS2, AS3, AS4, AS5 and AS6), and residual (E1, E2, E3, E4, E5 and E6) factors. Common (ac) and specific (as) genetic factor loadings and residual factor loadings (e) are given with respect to these factors, as shown in Supplementary Figure 2. For residual factor loadings (e) the first index points to the direction of the effect (i.e. the trait) and the second to the origin of the effect (i.e. the factor). 95% confidence intervals, assuming normality, can be approximated as 1.96 * SE.

GSEM, Genetic-relationship matrix structural equation modeling; IPC, Combined Independent Pathway (genetic part) / Cholesky (residual part) model

#### Supplementary Table 7: Factorial co-heritabilities for reading fluency

| **Label per factor and trait** | **Estimate (SE)** |
| --- | --- |
| (ac1)^2^/SNP-h_(1)_^2^ | 1 (<0.01) |
| (ac2)^2^/SNP-h_(2)_^2^ | 0.91 (0.07) |
| (ac3)^2^/SNP-h_(3)_^2^ | 0.91 (0.05) |
| (ac4)^2^/SNP-h_(4)_^2^ | 0.82 (0.09) |
| (ac5)^2^/SNP-h_(5)_^2^ | 0.93 (0.04) |
| (ac6)^2^/SNP-h_(6)_^2^ | 0.92 (0.07) |
| (as1)^2^/SNP-h_(1)_^2^ | <0.01 (<0.01) |
| (as2)^2^/SNP-h_(2)_^2^ | 0.09 (0.07) |
| (as3)^2^/SNP-h_(3)_^2^ | 0.09 (0.05) |
| (as4)^2^/SNP-h_(4)_^2^ | 0.18 (0.09) |
| (as5)^2^/SNP-h_(5)_^2^ | 0.07 (0.04) |
| (as6)^2^/SNP-h_(6)_^2^ | 0.08 (0.07) |

Based on the GSEM IPC (best-fitting) model for reading fluency (Figure 1, Supplementary Table 6, N=5,866). The model consists of (in that order of traits 1 to 6): NW reading a 9 (NBO), NW reading s 13 (TOWRE), W reading a 9 (NBO), W reading s 13 (TOWRE), P reading a 9 (NARA) and P reading s 9 (NARA). Factorial co-heritabilities reflect the proportion of genetic variance explained by a genetic factor with respect to the total SNP-h^2^ of the trait. Estimates are given here with respect to the squared factor loadings of the common genetic (ac1, ac2, ac3, ac4, ac5 and ac6) and the specific genetic (as1, as2, as3, as4, as5 and as6) factors. SEs were derived using the Delta method.

#### Supplementary Table 8: Genetic correlations, SNP-h^2^ and bivariate SNP-h^2^ for reading fluency

|  | NW reading a 9 | NW reading s 13 | | W reading a 9 | | W reading s 13 | P reading a 9 | P reading s 9 | |
| --- | --- | --- | --- | --- | --- | --- | --- | --- | --- |
| NW reading a 9 (NBO) | 0.30  (0.06) | | 0.61 (0.10) | 0.49 (0.08) | 0.62 (0.10) | | 0.53 (0.08) | | 0.59 (0.09) |
| NW reading s 13 (TOWRE) | 0.96 (0.04) | | 0.36  (0.07) | 0.67 (0.10) | 0.37 (0.08) | | 0.62 (0.09) | | 0.57 (0.09) |
| W reading a 9 (NBO) | 0.96 (0.03) | | 0.91 (0.04) | 0.45  (0.06) | 0.65 (0.10) | | 0.58 (0.07) | | 0.62 (0.08) |
| W reading s 13 (TOWRE) | 0.91 (0.05) | | 0.87 (0.05) | 0.87 (0.05) | 0.32  (0.08) | | 0.59 (0.10) | | 0.51 (0.10) |
| P reading a 9 (NARA) | 0.97 (0.02) | | 0.92 (0.04) | 0.92 (0.03) | 0.88 (0.05) | | 0.50  (0.06) | | 0.55 (0.08) |
| P reading s 9 (NARA) | 0.96 (0.04) | | 0.92 (0.05) | 0.92 (0.05) | 0.87 (0.06) | | 0.93 (0.04) | | 0.38  (0.07) |

Based on the GSEM IPC (best-fitting) model for reading fluency (Figure 1, Supplementary Table 6, N=5,866). The model consists of (in that order of traits 1 to 6): NW reading a 9 (NBO), NW reading s 13 (TOWRE), W reading a 9 (NBO), W reading s 13 (TOWRE), P reading a 9 (NARA) and P reading s 9 (NARA). Bivariate genetic correlations are shown in the lower triangle, bivariate SNP-h^2^ estimates in the upper triangle, and SNP-h^2^ estimates along the diagonal. Standard errors are shown in brackets. Bivariate SNP-h^2^ estimates reflect the proportion of the phenotypic covariance that is accounted for by the genetic covariance. Bivariate SNP-h^2^ SEs were approximated by the SE of the genetic covariance divided by the phenotypic covariance (as the SE of the phenotypic covariance is small).

#### Supplementary Table 9: Genetic and residual correlations for reading fluency

|  | NW reading a 9 | NW reading s 13 | | W reading a 9 | | W reading s 13 | P reading a 9 | P reading s 9 | |
| --- | --- | --- | --- | --- | --- | --- | --- | --- | --- |
| NW reading a 9 (NBO) | 1 | | 0.34 (0.05) | 0.60 (0.04) | 0.28 (0.05) | | 0.55 (0.04) | | 0.35 (0.05) |
| NW reading s 13 (TOWRE) | 0.96 (0.04) | | 1 | 0.37 (0.06) | 0.80 (0.03) | | 0.48 (0.06) | | 0.47 (0.05) |
| W reading a 9 (NBO) | 0.96 (0.03) | | 0.91 (0.04) | 1 | 0.34 (0.06) | | 0.61 (0.05) | | 0.41 (0.06) |
| W reading s 13 (TOWRE) | 0.91 (0.05) | | 0.87 (0.05) | 0.87 (0.05) | 1 | | 0.46 (0.07) | | 0.49 (0.05) |
| P reading a 9 (NARA) | 0.97 (0.02) | | 0.92 (0.04) | 0.92 (0.03) | 0.88 (0.05) | | 1 | | 0.59 (0.05) |
| P reading s 9 (NARA) | 0.96 (0.04) | | 0.92 (0.05) | 0.92 (0.05) | 0.87 (0.06) | | 0.93 (0.04) | | 1 |

Based on the GSEM IPC (best-fitting) model for reading fluency (Figure 1, Supplementary Table 6, N=5,866). The model consists of (in that order of traits 1 to 6): NW reading a 9 (NBO), NW reading s 13 (TOWRE), W reading a 9 (NBO), W reading s 13 (TOWRE), P reading a 9 (NARA) and P reading s 9 (NARA). Bivariate genetic and residual correlations are shown in the lower and upper triangle, respectively. Standard errors are shown in brackets.

#### Supplementary Table 10: Cholesky decomposition model (saturated model) for spelling, standardised factor loadings

| **Label** | **Path coefficient** | | **Variance explained (%)** |
| --- | --- | --- | --- |
|  | **Estimate (SE)** | **p** | **Estimate (SE)** |
| a11 | -0.59 (0.05) | 2.29 x 10^-33^ | 35.3 (0.06) |
| a21 | -0.55 (0.06) | 8.31 x 10^-24^ | 31.0 (0.06) |
| a22 | 0.14 (0.10) | 0.18 | 2.0 (0.03) |
| e11 | -0.81 (0.04) | 4.7 x 10^-114^ | 66.6 (0.06) |
| e21 | -0.55 (0.05) | 1.01 x 10^-33^ | 30.6 (0.05) |
| e22 | 0.61 (0.02) | 6.9 x 10^-147^ | 37.4 (0.03) |

Cholesky decomposition (saturated) model (Supplementary Figure 3, N=6,334). The model consists of (in that order of traits 1 to 2): Spelling a 7 (NB) and Spelling a 9 (NB). The phenotypic variance was dissected into genetic (A1, A2) and residual (E1, E2) factors. Genetic factor loadings (a) and residual factor loadings (e) are given with respect to these factors, such that the first index points to the direction of the effect (i.e. the trait) and the second to the origin of the effect (i.e. the factor).

#### Supplementary Table 11: Factorial co-heritabilities for spelling measures

| **Label per factor and trait** | **Estimate (SE)** |
| --- | --- |
| (a11)^2^/SNP-h_(1)_^2^ | 1.00 (<0.01) |
| (a21)^2^/SNP-h_(2)_^2^ | 0.94 (0.09) |
| (a22)^2^/SNP-h_(3)_^2^ | 0.06 (0.09) |

Factorial co-heritabilities based on the Cholesky decomposition (saturated) model for spelling measures (Supplementary Figure 3, Supplementary Table 10, N=6,334): The model consists of (in that order of traits 1 to 2): Spelling a 7 (NB) and Spelling a 9 (NB). Factorial co-heritabilities reflect the proportion of genetic variance explained by a genetic factor with respect to the total SNP-h^2^ of the trait. Estimates are given here with respect to the squared genetic factor loadings (a11, a21 and a22). SEs were derived using the Delta method.

#### Supplementary Table 12: Genetic correlations, SNP-h^2^ and bivariate SNP-h^2^ for spelling measures

|  | Spelling a 7 | Spelling a 9 | |
| --- | --- | --- | --- |
| Spelling a 7 (NB) | 0.35  (0.06) | | 0.46 (0.07) |
| Spelling a 9 (NB) | 0.97 (0.05) | | 0.33  (0.06) |

Cholesky decomposition (saturated) model for spelling measures (Supplementary Figure 3, Supplementary Table 10, N=6,334): The model consists of (in that order of traits 1 to 2): Spelling a 7 (NB) and Spelling a 9 (NB). Bivariate genetic correlations are shown in the lower triangle, bivariate SNP-h^2^ estimates in the upper triangle, and SNP-h^2^ estimates along the diagonal. Standard errors are shown in brackets. Bivariate SNP-h^2^ estimates reflect the proportion of the phenotypic covariance that is accounted for by the genetic covariance. Bivariate SNP-h^2^ SEs were approximated by the SE of the genetic covariance divided by the phenotypic covariance (as the SE of the phenotypic covariance is small).

#### Supplementary Table 13: GSEM IPC model for literacy-, language-, and PWM-related abilities (Data set: Passage reading), standardised factor loadings

| **Label** | **Path coefficient** | | **Variance explained (%)** |
| --- | --- | --- | --- |
|  | **Estimate (SE)** | **p** | **Estimate (SE)** |
| ac1 | 0.51 (0.05) | 3.51 x 10^-25^ | 27.0 (0.05) |
| ac2 | 0.56 (0.05) | 5.41 x 10^-29^ | 31.1 (0.06) |
| ac3 | 0.36 (0.06) | 2.80 x 10^-09^ | 13.3 (0.04) |
| ac4 | 0.37 (0.06) | 6.86 x 10^-11^ | 13.7 (0.04) |
| ac5 | 0.67 (0.05) | 7.11 x 10^-44^ | 45.0 (0.07) |
| as1 | 0.16 (0.08) | 0.05 | 2.6 (0.03) |
| as2 | 0.08 (0.19) | 0.67 | 0.7 (0.03) |
| as3 | -0.41 (0.08) | 1.35 x 10^-07^ | 16.6 (0.06) |
| as4 | -0.34 (0.07) | 3.11 x 10^-06^ | 11.9 (0.05) |
| as5 | 0.12 (0.15) | 0.42 | 1.4 (0.04) |
| e11 | 0.84 (0.03) | 7.32 x 10^-144^ | 73.6 (0.06) |
| e21 | 0.46 (0.04) | 1.22 x 10^-27^ | 21.0 (0.04) |
| e31 | 0.07 (0.04) | 0.13 | 0.4 (0.01) |
| e41 | 0.23 (0.04) | 1.34 x 10^-08^ | 5.3 (0.02) |
| e51 | 0.49 (0.04) | 5.84 x 10^-31^ | 24.5 (0.04) |
| e22 | -0.69 (0.02) | 9.39 x 10^-177^ | 47.6 (0.03) |
| e32 | 0.02 (0.03) | 0.57 | <0.01 (<0.01) |
| e42 | -0.13 (0.03) | 1.07 x 10^-05^ | 1.8 (0.01) |
| e52 | -0.09 (0.03) | 5.01 x 10^-03^ | 0.9 (0.01) |
| e33 | -0.84 (0.04) | 3.49 x 10^-108^ | 70.3 (0.06) |
| e43 | -0.12 (0.03) | 2.02 x 10^-05^ | 1.5 (0.01) |
| e53 | -0.04 (0.03) | 0.15 | 0.2 (<0.01) |
| e44 | -0.81 (0.03) | 1.03 x 10^-142^ | 66.0 (0.05) |
| e54 | -0.08 (0.02) | 6.94 x 10^-04^ | 0.7 (<0.01) |
| e55 | -0.53 (0.03) | 1.69 x 10^-85^ | 28.2 (0.03) |

GSEM IPC model for literacy-, language-, and PWM-related abilities (Data set: Passage reading; Figure 2, N=6,453). The model consists of (in that order of traits 1 to 5): Spelling a 7 (NB), Phon aware 7 (AAT), Listening compreh 8 (WOLD), NW repetition 8 (CNRep) and P reading s 9 (NARA). The phenotypic variance was dissected into common (AC1, AC2, AC3, AC4 and AC5), and specific genetic (AS1, AS2, AS3, AS4 and AS5), and residual (E1, E2, E3, E4 and E5) factors. Common (ac) and specific (as) genetic factor loadings and residual factor loadings (e) are given with respect to these factors, as shown in Supplementary Figure 4. For residual factor loadings (e) the first index points to the direction of the effect (i.e. the trait) and the second to the origin of the effect (i.e. the factor).

GSEM, Genetic-relationship matrix structural equation modeling; IPC, Combined Independent Pathway (genetic part) / Cholesky (residual part) model

#### Supplementary Table 14: Factorial co-heritabilities for literacy-, language-, and PWM-related abilities (Data set: passage reading)

| **Label** | **Estimate (SE)** |
| --- | --- |
| (ac1)^2^/SNP-h_(1)_^2^ | 0.91 (0.08) |
| (ac2)^2^/SNP-h_(2)_^2^ | 0.98 (0.10) |
| (ac3)^2^/SNP-h_(3)_^2^ | 0.44 (0.14) |
| (ac4)^2^/SNP-h_(4)_^2^ | 0.53 (0.14) |
| (ac5)^2^/SNP-h_(5)_^2^ | 0.97 (0.08) |
| (as1)^2^/SNP-h_(1)_^2^ | 0.09 (0.08) |
| (as2)^2^/SNP-h_(2)_^2^ | 0.02 (0.10) |
| (as3)^2^/SNP-h_(3)_^2^ | 0.56 (0.14) |
| (as4)^2^/SNP-h_(4)_^2^ | 0.47 (0.14) |
| (as5)^2^/SNP-h_(5)_^2^ | 0.03 (0.08) |

Based on the GSEM IPC model for literacy-, language-, and PWM-related abilities (Data set: Passage reading; Figure 2, Supplementary Table 13, N=6,453). The model consists of (in that order of traits 1 to 5): Spelling a 7 (NB), Phon aware 7 (AAT), Listening compreh 8 (WOLD), NW repetition 8 (CNRep) and P reading a 9 (NARA). Factorial co-heritabilities reflect the proportion of genetic variance explained by a genetic factor with respect to the total SNP-h^2^ of the trait. Estimates are given here with respect to the squared factor loadings of the five common (ac1, ac2, ac3, ac4 and ac5) and specific (as1, as2, as3, as4 and as5) genetic factor loadings. SEs were derived using the Delta method.

#### Supplementary Table 15: Genetic correlations, SNP-h^2^ and bivariate SNP-h^2^ for literacy-, language-, and PWM-related abilities (Data set: passage reading)

|  | Spelling a 7 | Phon aware 7 | Listening compreh 8 | NW repetition 8 | P reading a 9 |
| --- | --- | --- | --- | --- | --- |
| Spelling a 7 (NB) | 0.29  (0.06) | 0.45 (0.07) | 0.89 (0.18) | 0.56 (0.11) | 0.49 (0.07) |
| Phon aware 7 (AAT) | 0.94 (0.06) | 0.32  (0.06) | 1.00 (0.19) | 0.54 (0.11) | 0.58 (0.07) |
| Listening compreh 8 (WOLD) | 0.64 (0.11) | 0.66 (0.11) | 0.30  (0.06) | 0.55 (0.13) | 0.79 (0.15) |
| NW repetition 8 (CNRep) | 0.70 (0.10) | 0.72 (0.10) | 0.49 (0.10) | 0.26  (0.06) | 0.56 (0.10) |
| P reading a 9 (NARA) | 0.94 (0.05) | 0.97 (0.06) | 0.66 (0.11) | 0.72 (0.10) | 0.46  (0.06) |

Based on the GSEM IPC model for literacy-, language-, and PWM-related abilities (Data set: Passage reading; Figure 2, Supplementary Table 13, N=6,453). The model consists of (in that order of traits 1 to 5): Spelling a 7 (NB), Phon aware 7 (AAT), Listening compreh 8 (WOLD), NW repetition 8 (CNRep) and P reading a 9 (NARA). Bivariate genetic correlations are shown in the lower triangle, bivariate SNP-h^2^ estimates in the upper triangle, and SNP-h^2^ estimates along the diagonal. Standard errors are in brackets. Bivariate SNP-h^2^ estimates reflect the proportion of the phenotypic covariance that is accounted for by the genetic covariance. Bivariate SNP-h^2^ SEs were approximated by the SE of the genetic covariance divided by the phenotypic covariance (as the SE of the phenotypic covariance is small).

#### Supplementary Table 16: Genetic and residual correlations for literacy-, language-, and PWM-related abilities (Data set: passage reading)

|  | Spelling a 7 | Phon aware 7 | Listening compreh 8 | NW repetition 8 | P reading a 9 |
| --- | --- | --- | --- | --- | --- |
| Spelling a 7 (NB) | 1 | 0.55 (0.04) | 0.08 (0.05) | 0.27 (0.04) | 0.67 (0.03) |
| Phon aware 7 (AAT) | 0.94 (0.06) | 1 | 0.02 (0.05) | 0.27 (0.05) | 0.48 (0.05) |
| Listening compreh 8 (WOLD) | 0.64 (0.11) | 0.66 (0.11) | 1 | 0.16 (0.04) | 0.11 (0.07) |
| NW repetition 8 (CNRep) | 0.70 (0.10) | 0.72 (0.10) | 0.49 (0.10) | 1 | 0.31 (0.06) |
| P reading a 9 (NARA) | 0.94 (0.05) | 0.97 (0.06) | 0.66 (0.11) | 0.72 (0.10) | 1 |

Based on the GSEM IPC model for literacy-, language-, and PWM-related abilities (Data set: Passage reading; Figure 2, Supplementary Table 13, N=6,453). The model consists of (in that order of traits 1 to 5): Spelling a 7 (NB), Phon aware 7 (AAT), Listening compreh 8 (WOLD), NW repetition 8 (CNRep) and P reading a 9 (NARA). Bivariate genetic and residual correlations are shown in the lower and upper triangle respectively. Standard errors are in brackets.

#### Supplementary Table 17: GSEM IPC model for literacy-, language-, and PWM-related abilities (Data set: word reading), standardised factor loadings

| **Label** | **Path coefficient** | | **Variance explained (%)** |
| --- | --- | --- | --- |
|  | **Estimate (SE)** | **p** | **Estimate (SE)** |
| ac1 | -0.54 (0.05) | 3.73 x 10^-23^ | 30.1 (0.06) |
| ac2 | -0.58 (0.06) | 1.13 x 10^-23^ | 33.8 (0.07) |
| ac3 | -0.38 (0.07) | 5.22 x 10^-08^ | 14.3 (0.05) |
| ac4 | -0.35 (0.06) | 1.87 x 10^-08^ | 12.0 (0.04) |
| ac5 | -0.45 (0.07) | 3.20 x 10^-11^ | 20.6 (0.06) |
| as1 | -0.14 (0.14) | 0.32 | 2.0 (0.04) |
| as2 | 0.07 (0.32) | 0.83 | 0.5 (0.04) |
| as3 | -0.39 (0.09) | 6.78 x 10^-06^ | 15.0 (0.07) |
| as4 | 0.38 (0.07) | 3.75 x 10^-08^ | 14.4 (0.05) |
| as5 | -0.40 (0.08) | 1.05 x 10^-06^ | 16.5 (0.07) |
| e11 | 0.83 (0.04) | 4.06 x 10^-119^ | 70.6 (0.06) |
| e21 | 0.43 (0.05) | 1.96 x 10^-19^ | 18.4 (0.04) |
| e31 | 0.04 (0.05) | 0.42 | 0.2 (<0.01) |
| e41 | 0.23 (0.05) | 3.33 x 10^-07^ | 5.5 (0.02) |
| e51 | 0.31 (0.05) | 1.46 x 10^-09^ | 10.0 (0.03) |
| e22 | 0.69 (0.03) | 1.16 x 10^-162^ | 47.8 (0.04) |
| e32 | -0.03 (0.04) | 0.51 | 0.1 (<0.01) |
| e42 | 0.15 (0.04) | 5.20 x 10^-05^ | 2.2 (0.01) |
| e52 | 0.05 (0.04) | 0.20 | 0.3 (<0.01) |
| e33 | -0.84 (0.04) | 4.19 x 10^-108^ | 71.1 (0.06) |
| e43 | -0.13 (0.03) | 1.48 x 10^-05^ | 1.8 (0.01) |
| e53 | -0.08 (0.04) | 0.03 | 0.7 (0.01) |
| e44 | -0.80 (0.03) | 2.63 x 10^-133^ | 64.3 (0.05) |
| e54 | -0.09 (0.02) | 4.77 x 10^-05^ | 0.8 (<0.01) |
| e55 | 0.72 (0.04) | 8.87 x 10^-58^ | 52.4 (0.06) |

GSEM IPC (best-fitting) model for literacy-, language-, and PWM-related abilities (Data set: word reading accuracy, Figure 3, N=6,383). The model consists of (in that order of traits 1 to 5): of Spelling a 7 (NB), Phon aware 7 (AAT), Listening compreh 8 (WOLD), NW repetition 8 (CNRep) and W reading s 13 (TOWRE). The phenotypic variance was dissected into common (AC1, AC2, AC3, AC4 and AC5), and specific genetic (AS1, AS2, AS3, AS4 and AS5), and residual (E1, E2, E3, E4 and E5) factors. Common (ac) and specific (as) genetic factor loadings and residual factor loadings (e) are given with respect to these factors, as shown in Supplementary Figure 4. For residual factor loadings (e) the first index points to the direction of the effect (i.e. the trait) and the second to the origin of the effect (i.e. the factor).

GSEM, Genetic-relationship matrix structural equation modeling; IPC, Combined Independent Pathway (genetic part) / Cholesky (residual part) model

#### Supplementary Table 18: Factorial co-heritabilities for literacy-, language-, and PWM-related abilities (Data set: word reading)

| **Label** | **Estimate (SE)** |
| --- | --- |
| (ac1)^2^/SNP-h_(1)_^2^ | 0.94 (0.12) |
| (ac2)^2^/SNP-h_(2)_^2^ | 0.99 (0.13) |
| (ac3)^2^/SNP-h_(3)_^2^ | 0.49 (0.17) |
| (ac4)^2^/SNP-h_(4)_^2^ | 0.45 (0.13) |
| (ac5)^2^/SNP-h_(5)_^2^ | 0.56 (0.14) |
| (as1)^2^/SNP-h_(1)_^2^ | 0.06 (0.12) |
| (as2)^2^/SNP-h_(2)_^2^ | 0.01 (0.13) |
| (as3)^2^/SNP-h_(3)_^2^ | 0.51 (0.17) |
| (as4)^2^/SNP-h_(4)_^2^ | 0.55 (0.13) |
| (as5)^2^/SNP-h_(5)_^2^ | 0.44 (0.14) |

Based on the GSEM IPC model for literacy-, language-, and PWM-related abilities (Data set: word reading accuracy, Figure 3, Supplementary Table 17, N=6,383). The model consists of (in that order of traits 1 to 5): Spelling a 7 (NB), Phon aware 7 (AAT), Listening compreh 8 (WOLD), NW repetition 8 (CNRep) and W reading s 13 (TOWRE). Factorial co-heritabilities reflect the proportion of genetic variance explained by a genetic factor with respect to the total SNP-h^2^ of the trait. Estimates are given here with respect to the squared factor loadings of the common genetic (ac1, ac2, ac3, ac4 and ac5) and the specific genetic (as1, as2, as3, as4 and as5) factors. SEs were derived using the Delta method.

#### Supplementary Table 19: Genetic correlations, SNP-h^2^ and bivariate SNP-h^2^ for literacy-, language-, and PWM-related abilities (Data set: word reading)

|  | Spelling a 7 (NB) | Phon aware 7 (AAT) | Listening compreh 8 (WOLD) | NW repetition 8 (CNRep) | W reading s 13 (TOWRE) |
| --- | --- | --- | --- | --- | --- |
| Spelling a 7 (NB) | 0.32  (0.06) | 0.50 (0.08) | 0.98 (0.20) | 0.56 (0.12) | 0.54 (0.11) |
| Phon aware 7 (AAT) | 0.96 (0.06) | 0.34  (0.06) | 1.00* (0.21) | 0.53 (0.12) | 0.65 (0.12) |
| Listening compreh 8 (WOLD) | 0.68 (0.13) | 0.69 (0.12) | 0.29  (0.07) | 0.53 (0.14) | 0.69 (0.17) |
| NW repetition 8 (CNRep) | 0.65 (0.11) | 0.67 (0.11) | 0.47 (0.11) | 0.26  (0.06) | 0.52 (0.12) |
| W reading s 13 (TOWRE) | 0.72 (0.10) | 0.74 (0.10) | 0.52 (0.11) | 0.50 (0.10) | 0.37  (0.08) |

Based on the GSEM IPC model for literacy-, language-, and PWM-related abilities (Data set: word reading accuracy, Figure 3, Supplementary Table 17, N=6,383). The model consists of (in that order of traits 1 to 5): Spelling a 7 (NB), Phon aware 7 (AAT), Listening compreh 8 (WOLD), NW repetition 8 (CNRep) and W reading s 13 (TOWRE). Bivariate genetic correlations are shown in the lower triangle, bivariate SNP-h^2^ estimates in the upper triangle, and SNP-h^2^ estimates along the diagonal. Bivariate SNP-h^2^ estimates reflect the proportion of the phenotypic covariance that is accounted for by the genetic covariance. Bivariate SNP-h^2^ SEs were approximated by the SE of the genetic covariance divided by the phenotypic covariance (as the SE of the phenotypic covariance is small). *Truncated at 1.

#### Supplementary Table 20: Genetic and residual correlations for literacy-, language-, and PWM-related abilities (Data set: word reading)

|  | Spelling a 7 (NB) | Phon aware 7 (AAT) | Listening compreh 8 (WOLD) | NW repetition 8 (CNRep) | W reading s 13 (TOWRE) |
| --- | --- | --- | --- | --- | --- |
| Spelling a 7 (NB) | 1 | 0.53 (0.04) | 0.05 (0.06) | 0.27 (0.05) | 0.39 (0.06) |
| Phon aware 7 (AAT) | 0.96 (0.06) | 1 | -4x10^-4^ (0.06) | 0.29 (0.05) | 0.27 (0.06) |
| Listening compreh 8 (WOLD) | 0.68 (0.13) | 0.69 (0.12) | 1 | 0.16 (0.05) | 0.12 (0.06) |
| NW repetition 8 (CNRep) | 0.65 (0.11) | 0.67 (0.11) | 0.47 (0.11) | 1 | 0.24 (0.05) |
| W reading s 13 (TOWRE) | 0.72 (0.10) | 0.74 (0.10) | 0.52 (0.11) | 0.50 (0.10) | 1 |

Based on the GSEM IPC model for literacy-, language-, and PWM-related abilities (Data set: word reading accuracy, Figure 3, Supplementary Table 17, N=6,383). The model consists of (in that order of traits 1 to 5): Spelling a 7 (NB), Phon aware 7 (AAT), Listening compreh 8 (WOLD), NW repetition 8 (CNRep) and W reading s 13 (TOWRE). Bivariate genetic and residual correlations are shown in the lower and upper triangle. Standard errors are in brackets.

### Supplementary Figures

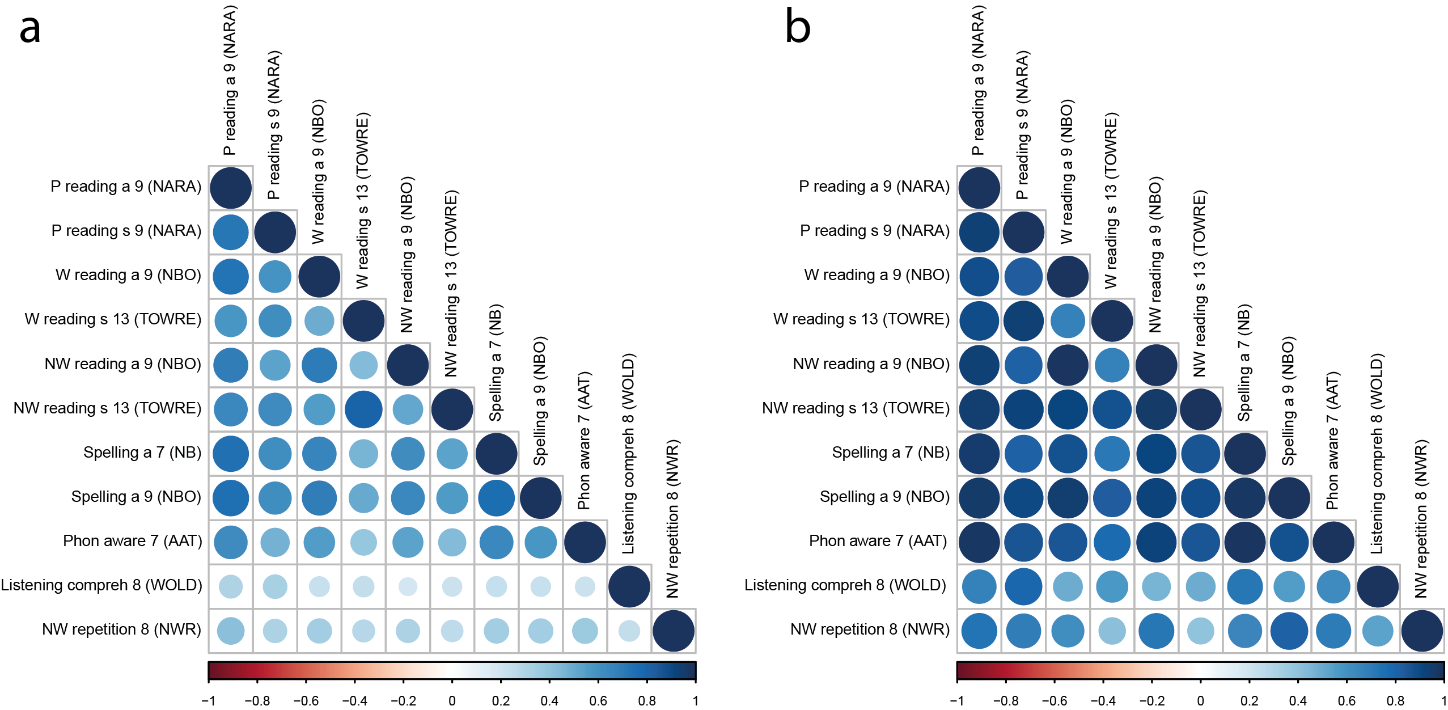

#### Supplementary Figure 1: Phenotypic and genetic correlations between literacy-, language-, and PWM-related abilities

Phenotypic correlations (a) were estimated with Pearson’s correlation coefficients (Upper triangle, Supplementary Table 2). Genetic correlations (b) were estimated with GCTA software (Supplementary Table 3). WORD, Wechsler Objective Reading Dimension; ALSPAC, Avon Longitudinal study of Parents and Children; NBO, ALSPAC-specific assessment developed by Nunes, Bryant and Olson; NARA II, The Neale Analysis of Reading Ability-Second Revised British Edition; TOWRE, Test Of Word Reading Efficiency; NB, ALSPAC-specific assessment developed by Nunes and Bryant; AAT, Auditory Analysis Test; WOLD, Wechsler Objective Language Dimensions; CNRep, Children’s Test of Nonword Repetition. All analyses are based rank-transformed residualized scores, using pairwise complete observations.

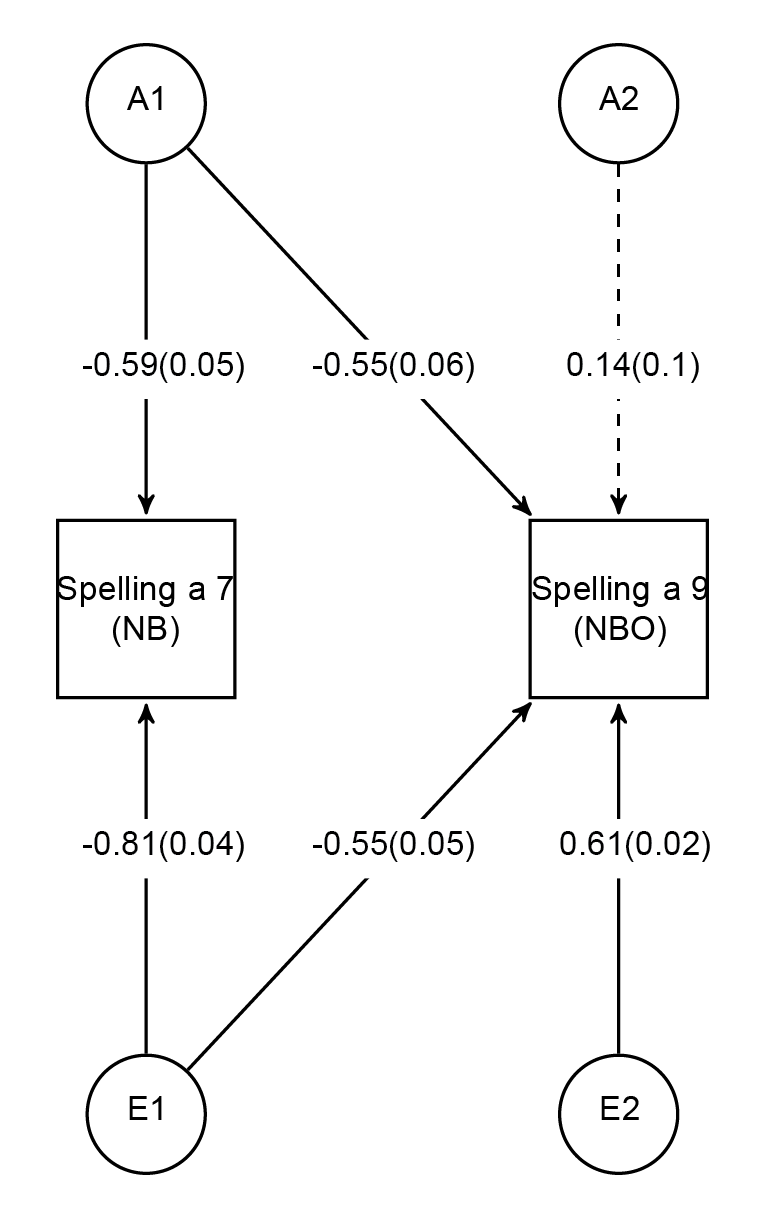

#### Supplementary Figure 2: Structural model of spelling

Genetic-relationship matrix structural equation modeling (GSEM) of two spelling measures (N=6334). The path diagram depicts the Cholesky model, here equivalent to an independent pathway model), for two measures of spelling accuracy at age 7 (Spelling a 7, NB) and age 9 (Spelling a 9, NBO). The phenotypic variance was dissected into genetic (A1 and A2) and residual (E1 and E2) factors.

Observed phenotypic measures are represented by squares, and latent factors are represented by circles. Single-headed arrows (paths) define relationships between variables. Dotted and solid paths represent factor loadings with p>0.05 and p≤0.05 respectively. Note that the variance of latent variables is constrained to unit variance; this is omitted from the diagram to improve clarity.

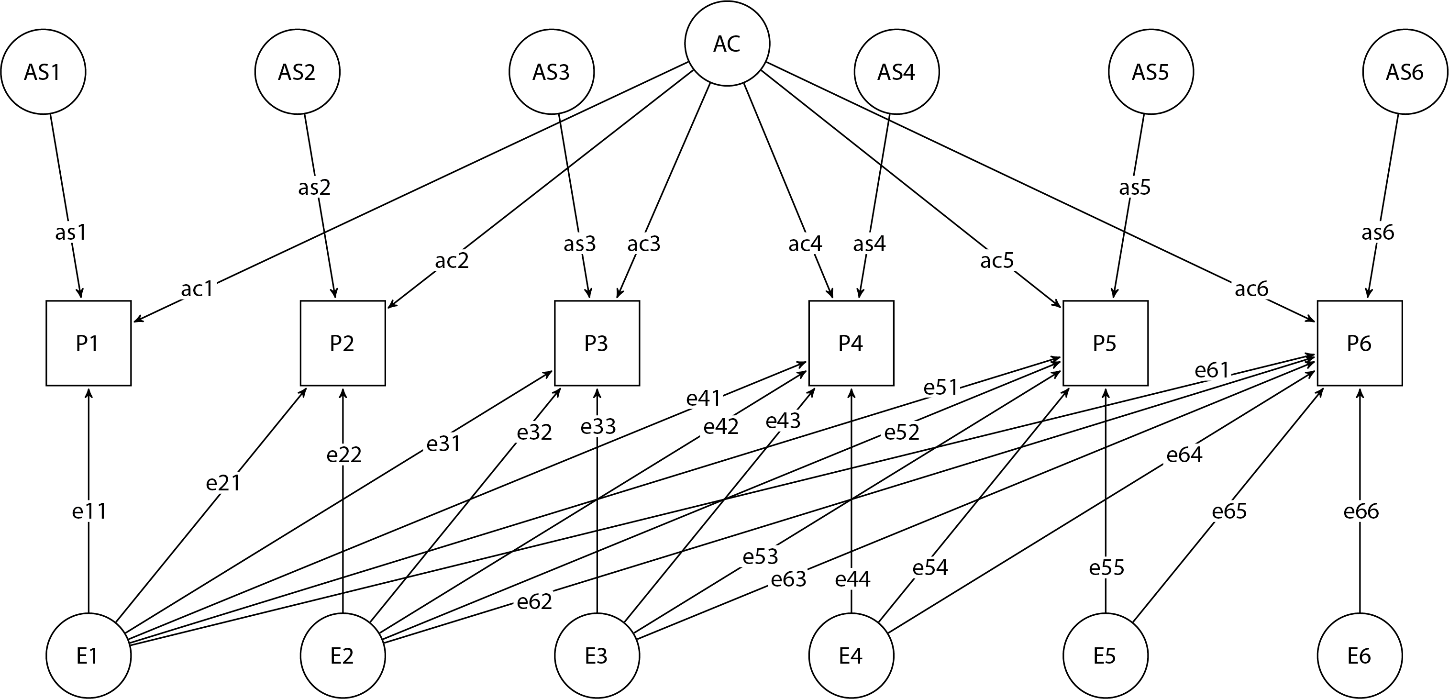

#### Supplementary Figure 3: Six-variate combined Independent Pathway / Cholesky model

Schematic diagram of a six-variate IPC model consisting of 6 phenotypes (P1, P2, P3, P4, P5 and P6). The phenotypic variance was dissected into latent common (AC) and specific (AS1, AS2, AS3, AS4, AS5 and AS6) genetic factors, as well as latent (E1, E2, E3, E4, E5 and E6) residual factors. Common (ac) and specific (as) genetic factor loadings and residual factor loadings (e) are given with respect to these factors. For residual factor loadings (e) the first index points to the direction of the effect (i.e. the trait) and the second to the origin of the effect (i.e. the factor). Observed phenotypic measures are represented by squares, and latent factors are represented by circles. Single-headed arrows (paths) define relationships between variables. Note that the variance of latent variables is constrained to unit variance; this is omitted from the diagram to improve clarity.

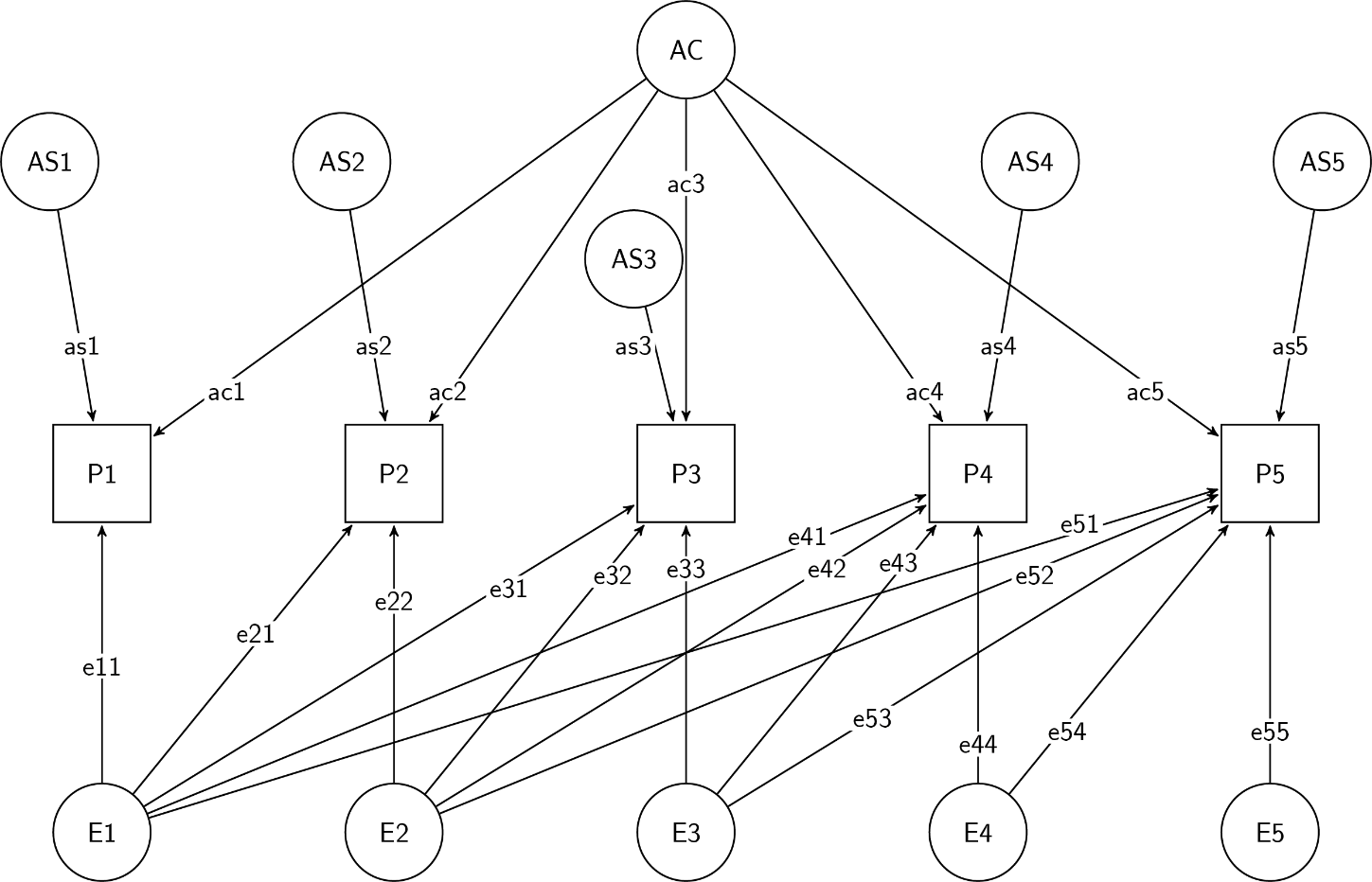

#### Supplementary Figure 4: Five-variate combined Independent Pathway / Cholesky model

Schematic diagram of a five-variate IPC model consisting of 5 phenotypes (P1, P2, P3, P4 and P5). The phenotypic variance was dissected into latent common (AC) and specific (AS1, AS2, AS3, AS4, AS5) genetic factors, as well as latent (E1, E2, E3, E4 and E5) residual factors. Common (ac) and specific (as) genetic factor loadings and residual factor loadings (e) are given with respect to these factors. For residual factor loadings (e) the first index points to the direction of the effect (i.e. the trait) and the second to the origin of the effect (i.e. the factor). Observed phenotypic measures are represented by squares, and latent factors are represented by circles. Single-headed arrows (paths) define relationships between variables. Note that the variance of latent variables is constrained to unit variance; this is omitted from the diagram to improve clarity.
